# Epistasis detection and fitness valleys in viral evolution

**DOI:** 10.1101/2025.09.05.674565

**Authors:** I.V. Likhachev, I.M. Rouzine

## Abstract

Probabilistic prognosis of virus evolution, vital for the design of effective vaccines and antiviral drugs, requires the knowledge of adaptive landscape including epistatic interactions. Although epistatic interactions can, in principle, be inferred from abundant sequencing data by various methods, it has been shown previously that genetic linkage between evolving sites obscures their signature and requires averaging over several independent populations. Here we probe the limits of epistasis detection based on pairwise correlations conditioned on the state of a third site on synthetic sequences evolved in a Monte Carlo algorithm with known epistatic pairs in two scenarios: an initially diverse population in the presence of recombination and mutation, with a uniform epistatic network, and an initially–uniform population in the presence of mutation and a fitness valley. In either scenario, the detection error decreases with the number of independent populations and increases with the sequence length. In the first scenario, the accuracy is enhanced by moderate recombination and is maximal, when epistasis magnitude approaches the point of full compensation. In the second scenario, simulation predicts the spontaneous crossing of a fitness valley and forming a new large strain.. The method is then applied to several thousands of sequences of SARS-CoV-2. Results obtained under equal sampling from world regions are consistent with the existence of fitness valleys connecting groups of viral variants.

**SIGNIFICANCE:** The few epistatic pairs of genomic sites hide in genomic data among numerous random correlations caused by common phylogenetic history. We test a method of epistasis detection designed to compensate for this noise. The accuracy is tested using synthetic sequences generated by a Monte Carlo algorithm with known epistatic pairs. The method is applied to several thousands of sequences of SARS-CoV-2 sampled in two different ways. Results obtained under equal sampling from world regions imply the existence of fitness valleys connecting groups of viral variants.

## INTRODUCTION

Elucidating the connection between genotype and phenotype remains a challenging problem. Most traits in organisms are coded by several genomic sites, whose combined effect on the phenotype is not a sum of their effects, but also includes site-site interaction termed “epistasis”. Epistasis is responsible for a significant fraction of genetic inheritance [1, 2]. In pathogens, epistasis facilitates the development of drug resistance and immune escape and impedes reversion of drug-resistant mutations [3–7]. Mutations in epitopes assisted by epistasis help a pathogen to escape the cytotoxic immune response in the host, when viral replication capacity lowered by an escape mutation is gradually restored later by compensatory mutations [8–11]. This process, termed fitness valley, is thought to be responsible for the high diversity and rapid HIV evolution in untreated patients [12].

Epistasis is also implicated in the evolution of variants of SARS-CoV-2 [13]. Some mutations in neutralizing antibody-binding regions that help SARS-CoV-2 to escape immune recognition during transmission [14–18] negatively affect the function of receptor binding, and have to be compensated elsewhere [19]. The tradeoff between antibody escape and transmissibility has been inferred in immunocompromised patients [20]. The same is likely true for mutations conferring escape from innate responses to SARS-CoV-2 [21, 22]

While pairwise epistasis can be measured from the free energy of binding [23] and in culture [24], these results cannot be transferred directly to the biological levels of an organism or population, where epistasis is defined with respect to fitness, i.e. the reproduction number at the within-host or between-host levels. Therefore, a large number of approaches has been proposed to measure epistasis directly from genomic data [25–27]. A popular method of “quasi-linkage equilibrium” assumes that recombination is rapid compared to allele frequency changes, and that population is large, so that recombination and the natural selection with epistasis are the dominant evolutionary factors, while the effects of genetic linkage are relatively small [28–31]. However, none of these methods enable reliable measurement of epistasis in asexual populations or in sexual populations at close loci [32].

About a century ago, it was realized that the evolution of a population is strongly affected by the fact that the fates of alleles at different loci are linked unless separated by recombination [33]. Empirically, the effect of strong linkage between genomic sites has been discovered by Thomas Hunt Morgan who used the inter-dependent segregation of alleles in fruit flies to understand the role of chromosomes and sex in inheritance. He showed that some pairs of traits of fruit flies segregate in a correlated fashion, and some in independent fashion. He postulated that the first are close on the same chain of genes, so that recombination cannot split them. Later, the nature of this chain was shown to be DNA in chromosomes.

Since then, a large body of experiment and modeling demonstrated that linkage changes evolutionary properties drastically. In the absence of recombination, common phylogeny makes all sequences very similar to their MRCA [34]. Moreover, even very frequent recombination does not break up linkage effects in a sufficiently long genome [35].

The increasingly dominant effect of linkage over epistasis in allelic associations originates from the genetic divergence of independent populations in time. All sequences in a population are similar to their most recent common ancestor, which moves away from the origin along a stochastic trajectory [34]. As a result, any measure of co-variance produces strong noise of random sign. An attempt at detecting epistasis, whether by using covariance measures, such as Pearson coefficient, mutual entropy, universal footprint [36], or by tree-based methods [37, 38] faces the same problem, the overwhelming genetic linkage effects. The only way to resolve this issue is to average the haplotype frequencies over many independent populations evolving under similar conditions. To decrease the required number of populations, it was proposed to use only sequences with a majority allele at a neighbor site of the measured site pair to interrupt a path along sites that creates a false-positive interaction [39].

In the present work, we test the fidelity of this detection technique using Monte-Carlo simulation in a broad range of parameters. Then, we apply the method to real virus sequences from the adapting world population of SARS–CoV-2 sampled in two different ways to predict the strongest epistatic links. This approach differs from the alternative methods in the following respects. Firstly, it does not make unrealistic assumptions regarding the evolutionary factors at play. Secondly, its accuracy is validated in a broad range of model parameters on simulated sequences, using a generic model of population genetics of the Wright Fisher type for two scenarios of evolution. We demonstrate the accuracy on simulated sequences under conditions resembling the evolution of SARS CoV-2 including the emergence of variants of concern. Thirdly, the application to data is focused on well-balanced sampling between sequences related to major variants. The fourth feature is the use of the conditional correlator.

## RESULTS

### Detection method

The method is based on the calculation of Universal Footprint correlator between pairs of loci [36]. This correlator, expressed in terms of haplotype frequencies by applying the second law of thermodynamics to a multi-locus evolutionary model, depends only on epistasis magnitude and its topology, but not on any other evolutionary parameters. The method takes a set of binarized sequences as initial data. We employ parallel computing on more than 10,000 processors of GPU to increase performance. To improve detection accuracy near the point of full compensation, an additional correlator conditioned on an allele at a third locus is calculated. Based on the relationship between the two correlators, an epistatic interaction between the two loci is inferred. The details are described in *Methods*.

### Accuracy test 1. The case of standing variation

To test the accuracy of the detection method, artificial genomic sequences are obtained from the simulation of the multi-site evolution of a genetically-diverse population by a Monte-Carlo algorithm with a set epistasis network (*Methods*). The input parameters are chosen in a range representative for several viruses (Table 1 in *Methods*). The predicted links are compared with the true links set in the algorithm. The error of detection is defined as the number of false-positive and false-negative interactions among predicted interactions divided by the number of true interactions. The error is calculated as a function of several parameters: genome length (total locus number), the number of independent populations used for averaging haplotype frequencies, the magnitude of epistasis, recombination probability, and the average crossover number (Fig. 1).

**Table 1.** Parameters of the model and data processing.

| Notation | Value | Description |
| --- | --- | --- |
| $f_0$ | 0 Figs. 2 to 4<br>0.45 Fig. 1 | Initial proportion of mutant alleles in the population |
| $N$ | 1000 | Population size |
| $\mu L$ | 0.07 Fig. 1<br>0.1 Figs. 2 to 4 | Mutation rate per genome |
| $t$ | 30 Fig. 1<br>300 Figs. 2 to 4 | Simulation time in generations |
| $s_0$ | 0.1 | Selection coefficient (the same for all sites) |
| $r$ | 0, 0.1, 0.5, or 1 | Recombination probability |
| $M$ | 1,3, or 10 | Number of crossovers |
| $E$ | 0.4,0.5,0.6,0.75 Fig. 1<br>0.4 - 4 Figs. 2 to 4 | Strength of epistatic interaction |
| $run$ | 6,20,50 or 200 | Number of simulation runs used to average the data |
| $L$ (seqlen) | 20,40,100 or 200 | Length of the simulated sequence |
| Sequence threshold | 0.7 | Maximum proportion of gaps and nonreads in one sequence |
| Gap threshold | 0.5 | Maximum proportion of gaps and nonreads at one site |
| Monomorphism threshold | 0.8 or 0.95 | Maximum degree of monomorphism per site |

**Figure 1.**
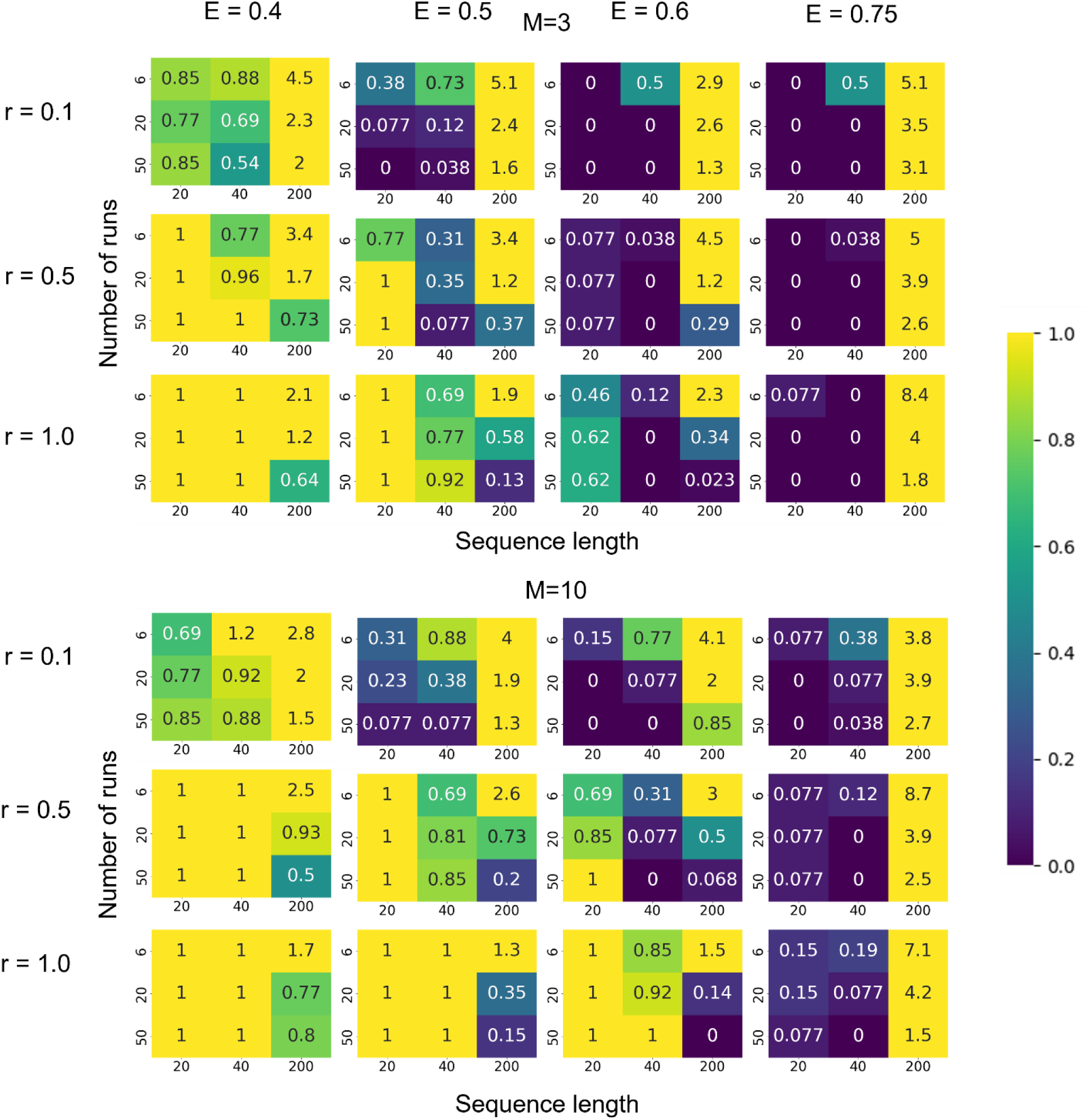
The error of detection of epistatic interactions for the case of standing variation (Test1). The length of the sequence is plotted on the X–axis, and the number of runs is plotted on the Y-axis of each panel. The values inside the squares defined by color map on the right correspond to the sum of the false positive and false negative interactions divided by the number of true interactions. Detection is carried out using the universal footprint of epistasis (UFE) [36] and three-site haplotype method [39] (*Methods*). All values greater than 1 are set to have the same red color as values equal to 1. Testing of the method was conducted on the sequences generated by Monte Carlo simulation (*Methods*) with variable parameters (shown): probability of recombination *r*, number of crossovers *M*, strength of the epistatic interaction *E*. The topology of the epistatic links (interactions) comprises isolated double arches, for which *E* = 0.75 is the threshold value of full compensation [36]. Fixed parameters: initial allele frequency *f*_0_ = 0.45, population size *N* = 1000, mutation rate per genome *μL* = 0.07, simulation time in generations *t* = 30, selection coefficient *s* = 0.1. Results for *M* = 1 are shown in Figure S1. Variant without recombination is shown in Figure S2. Results using Pearson coefficient, with and without recombination, are in Figs. S3 and S4.

As expected, the accuracy of epistasis detection increases with the number of independent populations used for averaging and the magnitude of epistasis, and decreases with the genome length, as long as recombination is infrequent or absent. While modest recombination significantly improves the accuracy of detection by compensating linkage effects eclipsing epistasis, very frequent recombination counteracts associations of alleles caused by epistasis and so impairs detection (Fig. 1). Analogous procedure was carried out using Pearson coefficient and UFE [36] in the absence of recombination (*Supplement,* Figs. S2-S4).

### Accuracy test 2. Initially-monomorphous populations with a fitness valley

In real world, not all populations are polymorphous in the beginning of evolution. Therefore, we carried out an additional simulation test with initial allelic frequency at each locus *f*_0_ = 0. The epistatic network was changed from the isolated double arcs to a single star comprising one deleterious allele and 10 rays with compensatory alleles. Recombination is absent.

Initially, the population is monomorphous. Then a gradual accumulation of beneficial alleles occurs, which are assumed to take a 10% of loci. Deleterious alleles also accumulate at the remaining 90% of loci due to the hitchhiking effect, i.e., genetic linkage with the beneficial alleles, although more slowly than beneficial alleles. Among deleterious alleles, one is set to be the primary mutation of fitness valley and 10 are set to be compensatory mutations with the degrees of compensation ranging from 80 to 800%. These mutations are deleterious separately, but the combination of the primary allele with the compensatory alleles is advantageous. Sooner or later, deleterious primary mutation becomes abundant in a population due to the hitchhiking effect. Then compensatory alleles become advantageous, and start rapid accumulation, leading to a jump in the curve (Fig. 2). This marks passing the fitness valley and the emergence of a new group of virus variants on the end of fitness valley.

**Figure 2.**
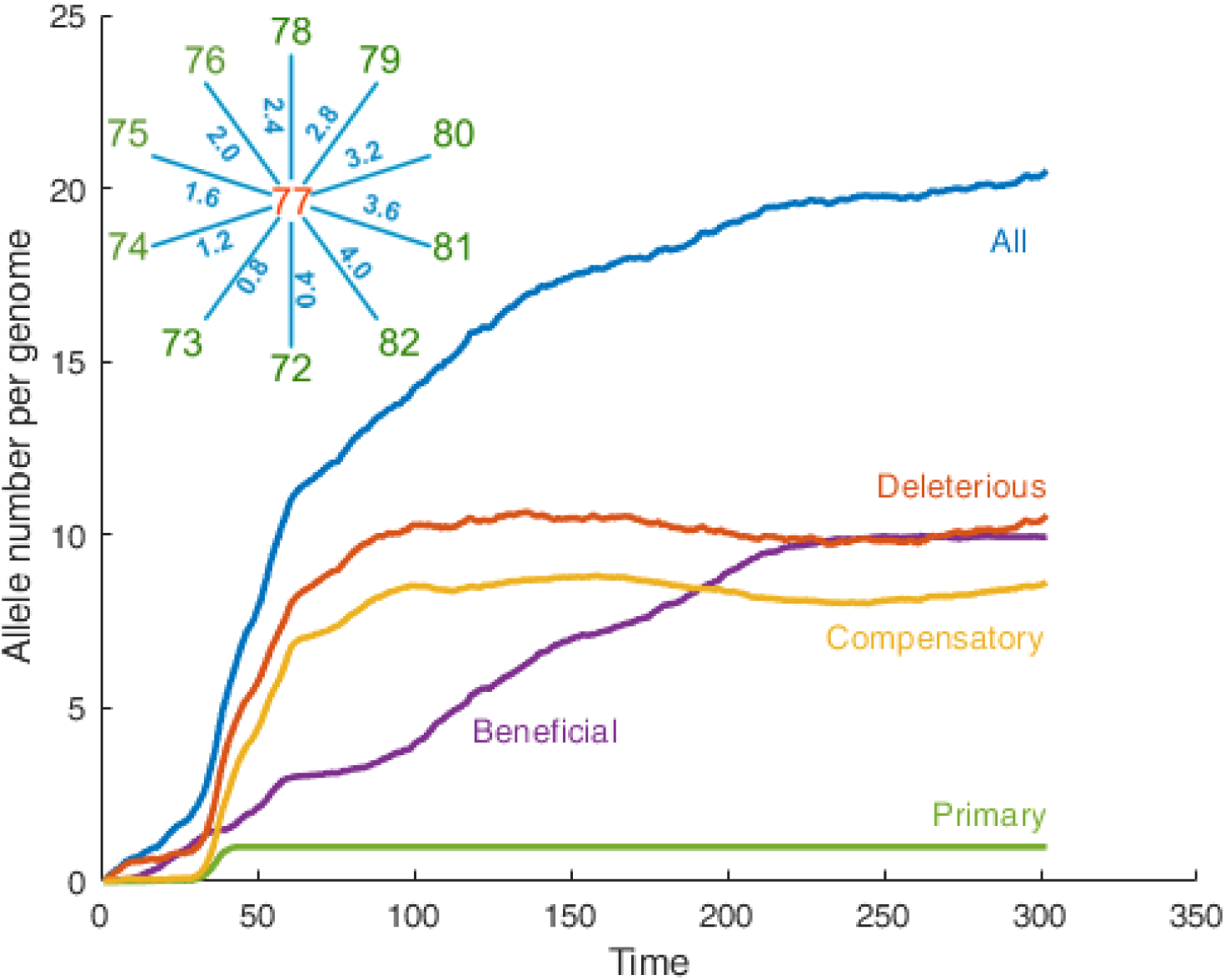
Monte Carlo simulation of the evolutionary dynamics of a population with a fitness valley. X-axis: Time in generations. Y axis: Average numbers of the alleles of different classes (shown) per genome. Parameters: initial allele frequency *f*_0_ = 0, population size *N* = 1000, mutation rate per genome *μL* = 0.1,10% of the alleles are beneficial, 90% are deleterious with selection coefficient *s* = 0.1 . Epistatic network represents a star with the primary allele in the center and ten compensatory mutations as the rays, all of which are deleterious as single alleles. Epistatic parameters are given in *Methods* Recombination is absent.

As a result of virus passing the fitness valley, the mutational spectrum averaged over 6 populations at 150 generational steps had two maxima (Fig. 3). The sequences were divided between the two peaks with borders chosen at 6 (the local minimum) and 22 new alleles (upper cutoff). Then we conducted bootstrapping procedure, in which sequences from each peak were oversampled many times with replacement and repetition. The resulting samples were used for the epistasis detection method described above.

**Figure 3.**
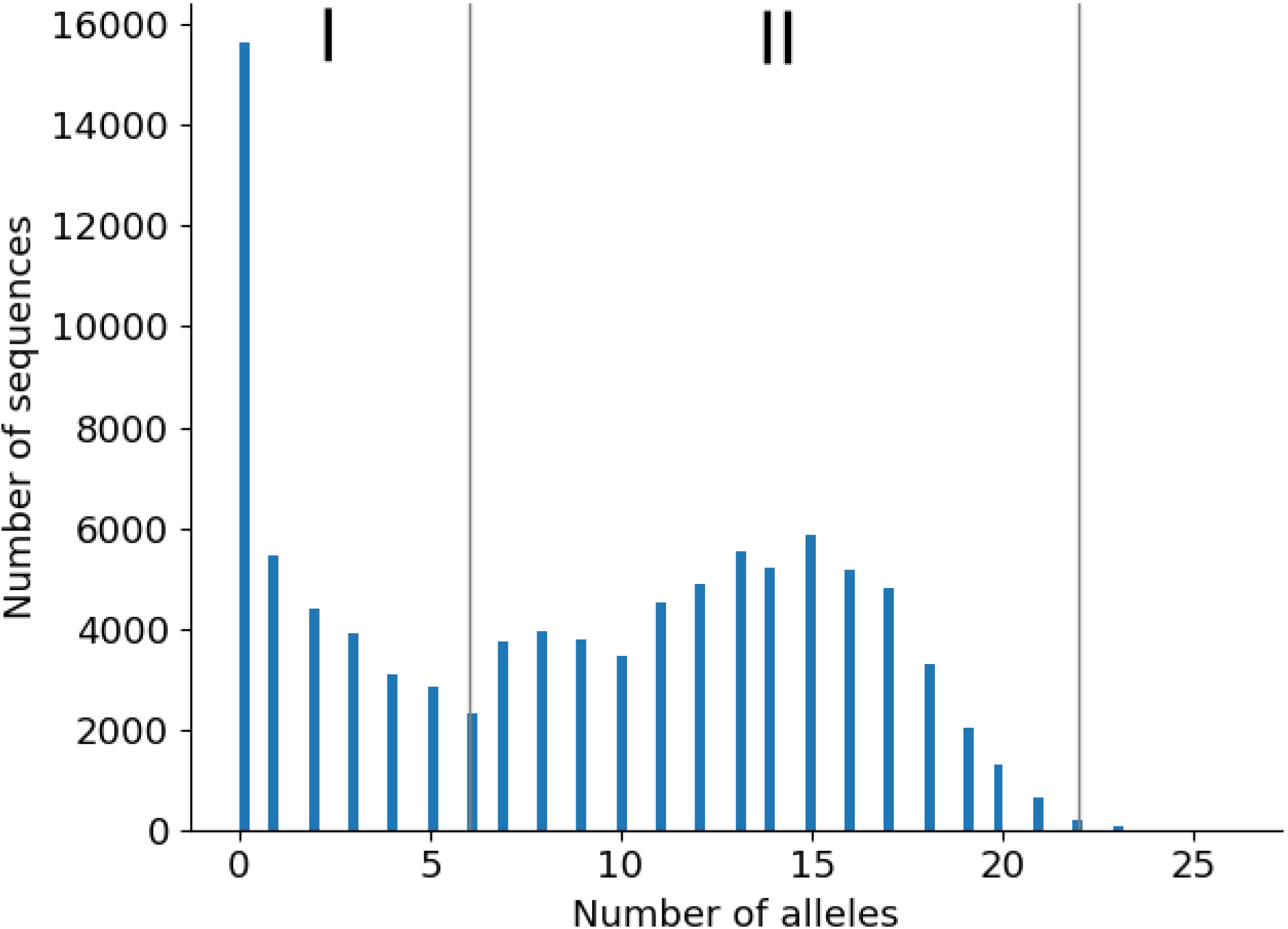
Mutational spectrum for six simulated populations, initial state is monomorphous. Y–axis: number of sequences with a number of replacements shown in X–axis. Gray lines at 6 and 22 replacements divide the two groups of sequences numbered by Roman numerals. Simulation parameters are as in Figure 2. Six Monte Carlo runs at timepoints multiple of 10 until time 150 are used for sampling.

The resulting detection error is calculated as the sum of false positive and false negative epistatic interactions out of 10 possible (Fig. 4), as a function of numbers of sequences sampled from peak 1 and 2, normalized to the sequence number of each peak. Thus, about a half of weaker interactions is not detected.

**Figure 4.**
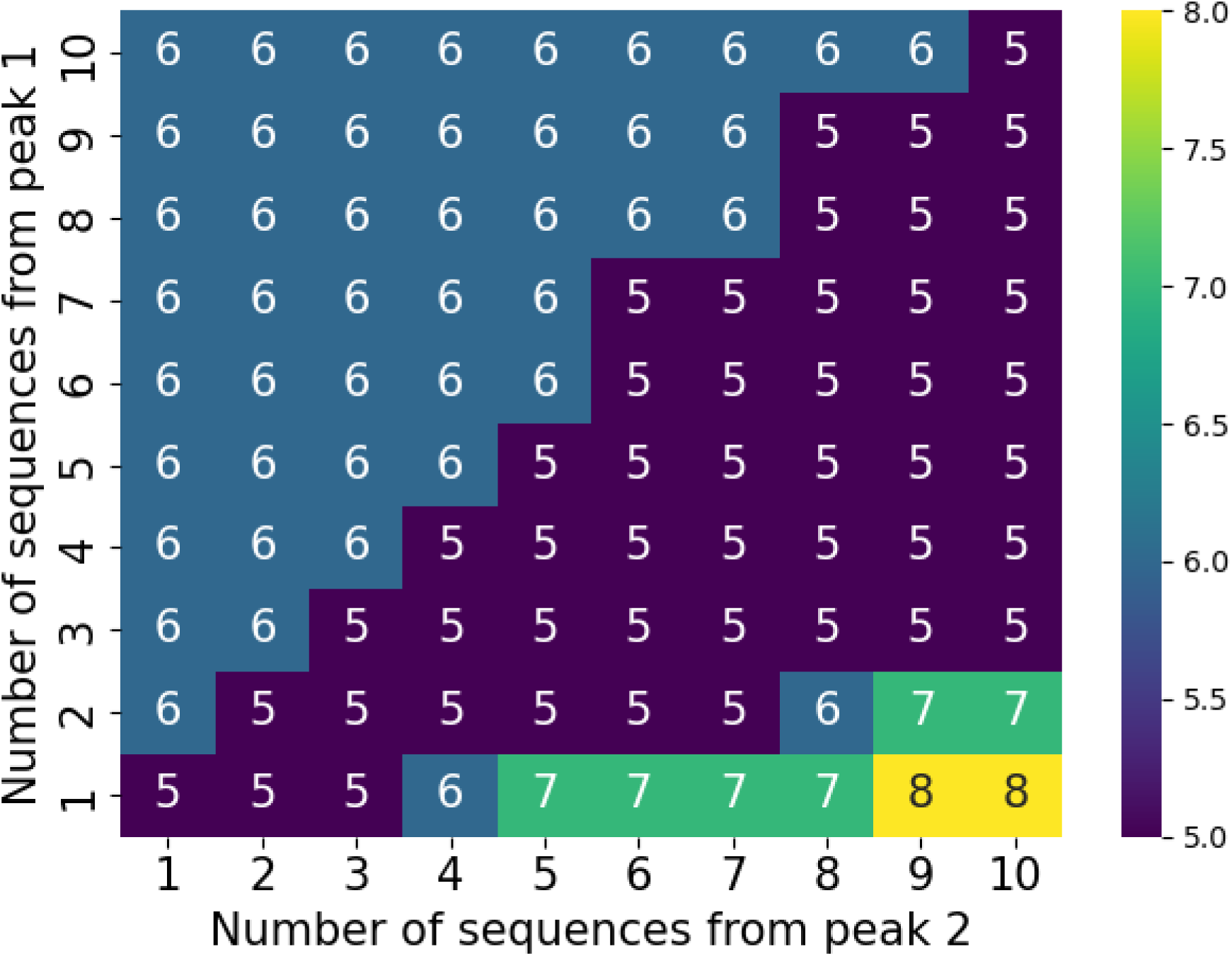
Error of epistatic detection for six simulated populations that are initially monomorphous. Axes X and Y show the rescaled numbers of sequences sampled from parts I and II of the dataset in Fig. 3, respectively, relative to the total number of sequences in each part. Numbers in cells show the number of false-negative epistatic interactions out of 10 possible. The measured number of false–positives is always zero. Parameters are as in Figure 2.

The number of false positive interactions remained zero, so that the total error was given by false negatives only . Therefore this method is highly accurate, but only in the conservative direction. Our results suggest the need to take either equal sampling from the two peaks or somewhat smaller sampling from the earlier peak (Peak I), as long as they are approximately equal in size, as in our example.

### Results on epistasis detection in genomic data for SARS CoV-2 are consistent with fitness valleys

After testing the method on simulated sequences, we applied it to sequence data on SARS-CoV-2 for two different data sets, as follows.

Dataset 1 includes 6000 sequences from 6 regions, 1000 from each: Africa, Asia, North America, Oceania, South America, for the period of one year, 09/24/2020 to 09/25/2021. When each sequence is compared to Wuhan variant, a mutation spectrum is obtained, where sequences segregate naturally into three groups, with gaps between them (Fig. 5). .

**Figure 5.**
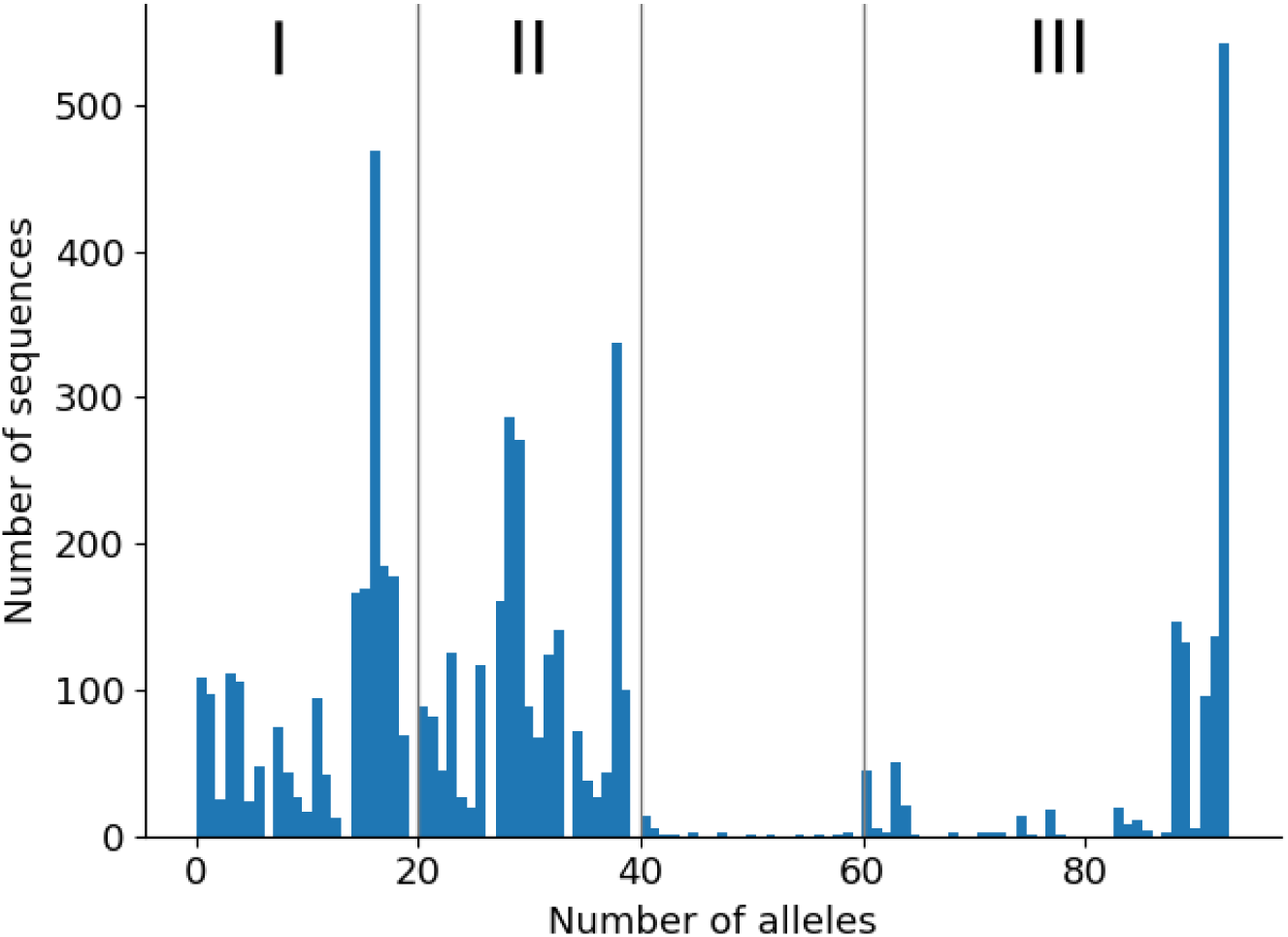
Mutational spectrum for Dataset 1. Y–axis: number of sequences with the number of differences shown in X–axis from the consensus sequence, defined as Wuhan variant. Gray lines at 20, 440 and 60 divide the three groups of sequences numbered by the Roman numerals.

Fig. 6 shows the inferred epistatic network from groups I and II. To obtain this diagram groups I and II were oversampled 10 times with replacement. As it was shown in Fig. 4 equal sampling is essential for obtaining reliable results. This procedure was also repeated for sequences mixed from groups I and III, where groups I and III were resampled 10 and 20 times respectively since group III almost two times smaller than group I (Fig. S5).

**Figure 6.**
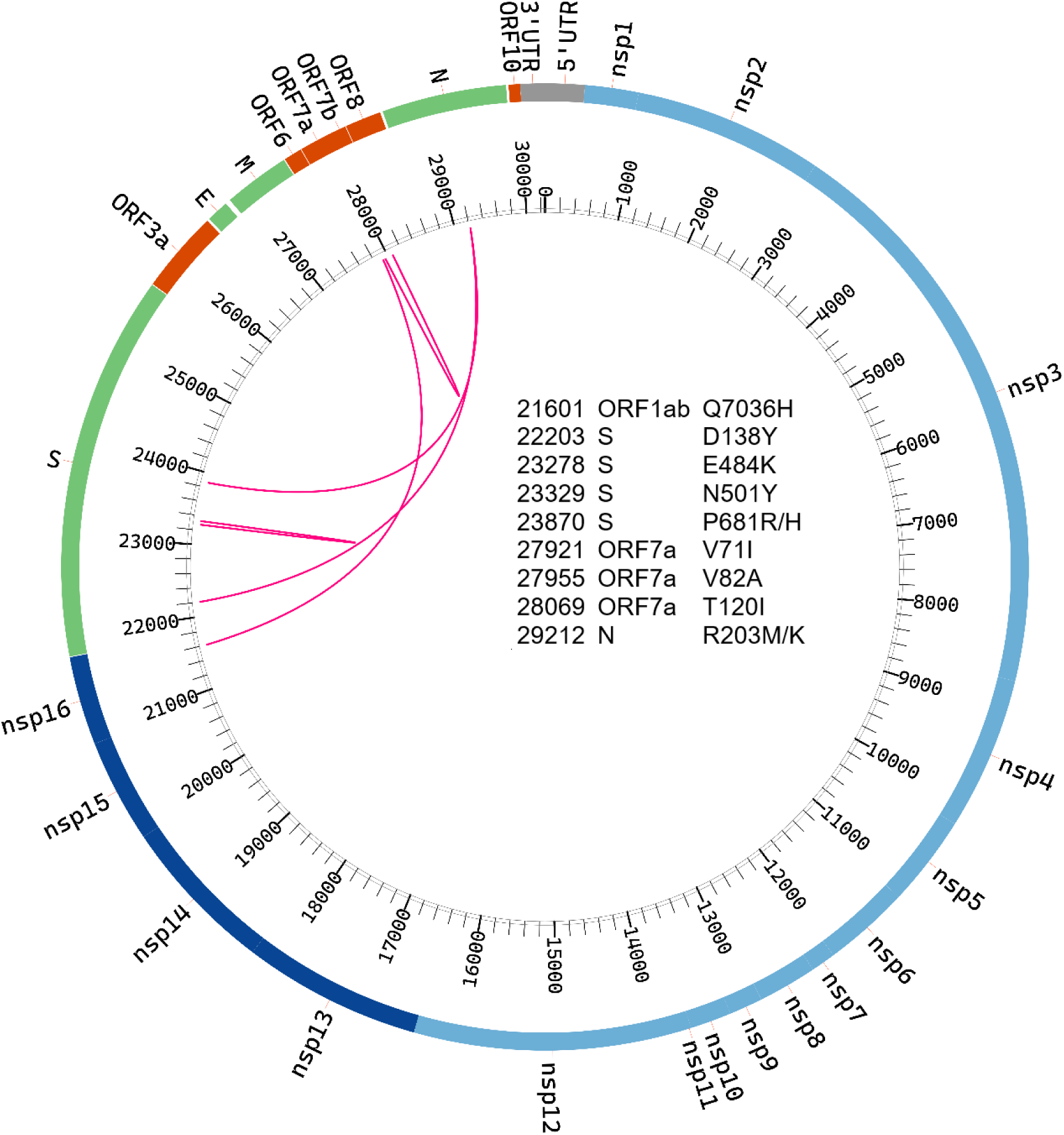
Epistatic interactions for SARS-CoV-2 genome predicted for Dataset 1 for sequence groups I and II (Fig. 5). To obtain the circular diagram using software Circos [40], each group was oversampled 10 times with replacement. The interacting pairs are shown.

The detected network (Fig. 6) is consistent with the presence of fitness valleys, although does not prove their existence directly, presumably, due to the absence of weaker links not captured by the method. The method is very conservative, and its accuracy at small epistatic magnitudes is low (Figs 1 and 4 and the legend to Fig. 4). Hence the epistastic interactions in Figs. 6 are the strongest in possible fitness valleys.

For sequences sampled from groups II and III (Fig. 5), no significant epistatic links are found, indicating the absence of a direct passage between them. The results we obtain are consistent with the existence of several fitness valleys connecting large groups of virus variants.

We repeated the same procedure for Dataset 2 comprising 20,000 random sequences from the NCBI Virus database for the same period of time as Dataset 1, from 09/24/2020 to 09/25/2021, but without geographic classification nor equal sampling from continents. Results differ strongly from those for Dataset 1, as follows. The number of interactions is smaller, and they are detected only for groups I and II, but not any other pairs of the four groups (Fig S6). Thus, equal sampling from world regions is critical for detection.

### Biological function of the predicted interacting sites

Mutations E484K and N501Y are well-known mutations of the receptor-binding domain of the coronavirus in Spike. The first is responsible for the viral escape from the immune response and is capable of reducing the affinity of antibodies produced to earlier strains by 1-2 orders of magnitude , while the second improves the binding of coronavirus to the human ACE2 receptor on cell surface and increases viral replication in upper respiratory tract cells. [41]

Mutations P681H and P681R, most often found in Alpha (B.1.1.7) and Delta (B.1.617.2) variants of the virus, respectively, are located in the furin cleavage region of Spike. They both confer a change in the amino acids from nonpolar hydrophobic to positively charge. They affect tropism (repertoire of infectable cells) by increasing the cleavage of S1/S2 proteins in human airway epithelial cells [42].

Mutation D138Y in Spike is believed to affect the protein’s secondary structure thus reducing antibody -binding efficiency [43]. The R203K mutation in the N protein was shown to increase transmission and virulence of SARS-COV-2 [44], while the R203M mutation, which creates a start codon for a new open reading frame, was reported to suppress interferon induction, thereby increasing viral infectivity [45].

V82A and T120I in ORF7a are most frequently found in variant Delta of the coronavirus [46]. They are located within the di-lysine motif of ORF7a, which is important for the localization of viral proteins in endoplasmic reticulum. Existing data on mutation Q7036H in protein ORF1ab and mutation V71I in ORF7a do not permit to determine their functions definitively; however, their location within viral non-structural proteins hint at their interaction with the immune system of the host.

## DISCUSSION

Detection and measurement of epistasis from genetic data represents a challenge. In our work, we have upgraded a linkage-resistant method of detection and applied it to sequences from SARS-CoV-2. Results are consistent with the existence of fitness valleys that provide passages between large groups of virus variants.

As mentioned before, a fitness valley comprises a transient decrease in fitness due to initial mutation(s) forced by changes in the environment, which is followed by compensatory mutations. The fitness valley effect was previously observed or inferred for various viruses, including HIV [47, 48] and influenza [37–39], and studied using mathematical models [8, 39]. For example, a primary mutation can help a virus to escape the immune response, adapt to a species or a tissue, or expand the pH range in which it can replicate. Mutations conferring these benefits to a virus, but reducing its overall fitness, are followed by compensatory mutations that elevate its fitness above the initial level.

The loss of fitness due to escape mutations, and the partial loss of recognition by the immune system, together determine the order, the number, and the evolution rates of escape mutations in HIV infection [49, 50]. Compensatory mutations prevent the reversion of escape mutations after the decay of CD8 T cell clones against escaped epitopes. Bacteria developing antibiotic resistance also pass through such fitness valleys [5]. A population-level simulation study proposed this effect in conjunction with the evolution in immunocompromised patients infected with SARS-CoV-2 [51]. Another modeling work proposed crossing a deep fitness valley using recombination [52].

The fates of alleles at different loci are linked due to common phylogeny unless separated by recombination [33]. For this reason, one-locus models usually do not apply in real populations, and multi-locus models must be used instead [53, 54]. These models, based on traveling-wave technique, have been successfully applied to rapidly-evolving viruses, such as HIV [55] and influenza [56–58]. Linkage effects include clonal interference [33, 59], background selection, genetic hitchhiking [60], enhanced accumulation of deleterious alleles (Muller’s ratchet) [61, 62], and the increase of genetic drift at one locus due to selection at another [63]. Multi-site adaptation has been analyzed analytically in various asexual models, with the result that linkage effects strongly decrease the adaptation rate compared to the limit of independently-evolving sites [62, 64–66]. Importantly, genetic linkage also creates random associations between pairs of mutations occurring on the same branch of the ancestral tree, which is a serious obstacle to the detection of epistatic interactions in genetic data [32]. In a single asexual population, stochastic linkage completely overshadows the epistatic footprint, except in a narrow range of times and parameters [34]. The same limitation exists for the tree-based methods of detection [37, 38].

For testing the accuracy, we used a model with directional selection. In fact, SARS-CoV–2 evolves in antibody–binding regions due to collective immunity of population, which is a case of the selection for change (Red Queen effect). Nevertheless, as it has been shown by several groups [56–58], the multi-locus epidemiological model of respiratory viruses is effectively reduced to a Wright-Fisher model with constant directional selection. The selection coefficient is expressed in terms of basic reproduction ratio in a naïve population and the breadth of antibody neutralization, calculated in these works using different approximations.

The most popular method for searching for epistasis signature in genomic data is the “quasi linkage equilibrium” approach [28, 30], which assumes frequent recombination compared to changes in allelic frequency and a large population size. In this limit, genetic linkage effects are negligible compared to epistasis and recombination. In the present work, we do not rely on QLE approximation, so that the method works for finite population size and for modest recombination or even in its absence. The linkage effects are mostly cancelled out by averaging over well-balanced independent populations and the use of 3-site haplotypes. Such cancellation was not carried out in [68], hence the abundance of false negative links (Fig. 7C) compared to our results (Fig. 7A).

**Figure 7.**
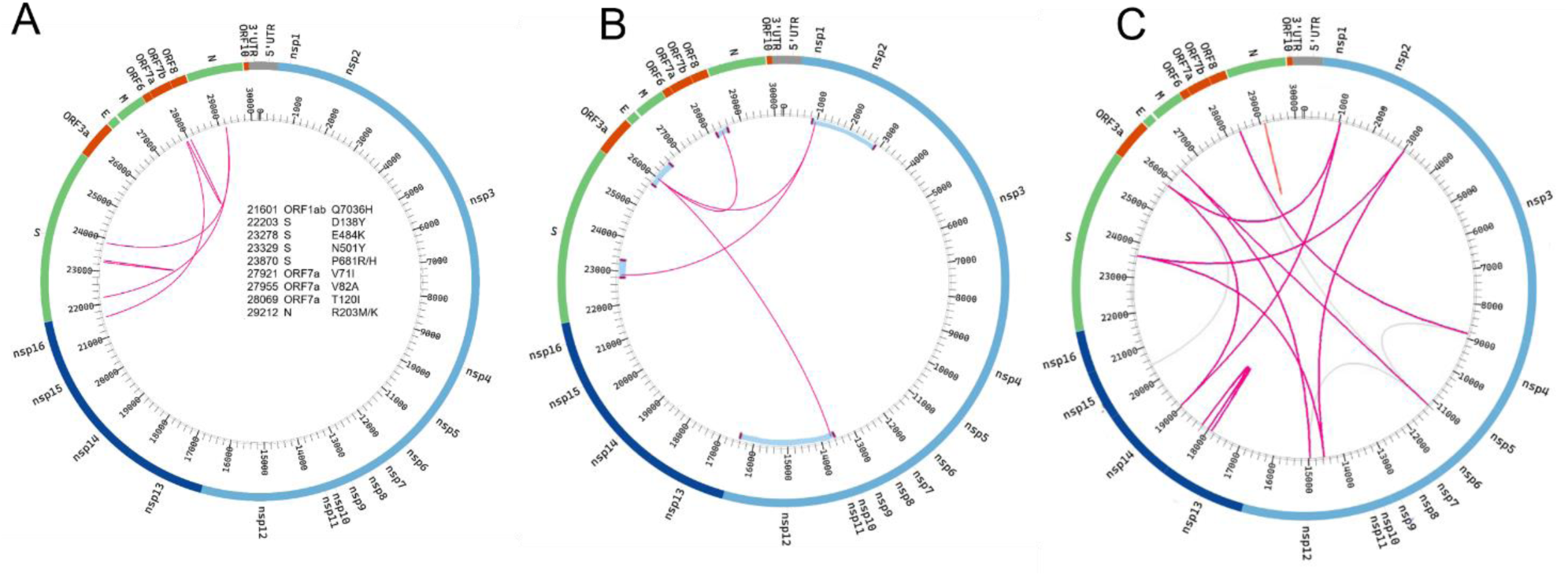
Epistatic interactions for SARS-CoV-2 genome: comparison of the present results with two previous studies. A) Results of the present work for Dataset 1 from Fig. 6 obtained from mixing sequence groups I and II in Fig. 5. B, C) Results from studies [67] and [68], respectively. Blue stripes in panel B represent protein domains reported to interact with each other in [67].

To address the problem of false-negative bonds, another team [67] compared the sequences of SARS CoV-2 not between themselves, but with other coronaviruses. They used “direct coupling analysis”, which postulates a statistical distribution of sequences (usually, of a Gaussian form) including individual effects of alleles and their pairwise interaction. A physicist would recognize here the Gibbs distribution for the Onsager model of ferromagnetism. In population genetics, this Gaussian form has a justification in the limit of QLE [28, 30]. Parameters of the Gaussian are fit to sequence data using various approximations [67]. DCA also includes down-weighing sequences with over 80% identity, which is a phenomenological trick to deal with the effects of random linkage. As result, the number of false-positive epistatic bonds due to genetic linkage is sharply decreased (Fig. 7B). The problem here is that the nature of such correction is not derived from a model of population genetics. QLE does not work in SARS CoV-2 evolution, because linkage effects, negligible in QLE limit, are quite strong.

Note that the use of tree-based methods does not automatically compensate for strong linkage effects. This is because, in the absence of sufficiently strong recombination, all sequences in one single population are very similar to their MRCA, and MRCAs for different independent populations are diverging from each other in genetic space. This process was demonstrated to be responsible for strong linkage effects [34]. To exclude the effects of linkage, one group used synonymous sites as markers of linkage [37, 38].

To conclude, the present study confirms the existence of fundamental limitations on epistasis detection due to genetic linkage. Within this limitation, a conditional correlation method combined with balanced sampling from quasi-independent populations and testing the accuracy in simulation allows the testable detection of strong epistatic links.

## METHODS

### Generating sequences for method testing

Evolution of a haploid asexual population of *N* binary sequences is simulated by a Monte Carlo code [36, 69], as follows. In an individual genome, each locus (site, nucleotide position, amino acid position) numbered *i* = 1, 2, … , *L* is occupied by one of two alleles, either the wild-type allele, denoted *K_i_* = 0, or the mutant allele, *K_i_* = 1.We use a discrete generation scheme in the absence of generation overlap (Wright-Fisher model). The evolutionary factors included in the model are random mutation with rate *μL* per genome, constant directional selection, random genetic drift due to random sampling of progeny and recombination with rate *r*. Selection includes an epistatic network with a set strength and topology. The logarithm of the average progeny number of a genome, *W*, depends on sequence [*K_i_*] as given by [36]

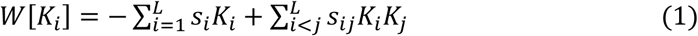

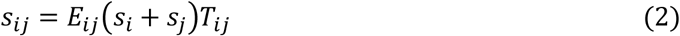

where *T_ij_* = 0 or 1 is the binary matrix that shows epistatically-interacting pairs. Here the selection coefficients *s_i_* and *s_j_* denote the individual fitness costs of two deleterious mutations, *E_ij_* is the compensation degree at sites *i* and *j*.

In our simulation example in Fig. 1, we consider a haploid population with the initial frequency of deleterious alleles *f*_0_ = 0.45, mutation rate per genome *μL* = 0.07, fixed selection coefficient *s_i_* ≡ *s* = 0.1, population size *N* = 1000 individuals, genome length *L* = 40 sites, and epistatic network is double arches

In our simulation example in Figs. 2 to 4, we consider a haploid population with the initial frequency of deleterious alleles *f*_0_ = 0, mutation rate per genome *μL* = 0.1, selection coefficients with a fixed absolute value *s_i_* ≡ *s* = 0.1, where 10% of potential mutations are advantageous and 90% are deleterious, population size *N* = 1000 individuals and genome length *L* = 100 sites, epistatic network – star with deleterious mutation in the middle and 10 compensatory mutations as rays, degree of compensation gradually increases from 80% to 800%

The Monte-Carlo simulation code is written in MATLAB^TM^ and deposited at https://github.com/FireFrog17/Fundamental-restriction-on-epistasis-detection-and-fitness-valleys-in-virus-evolution-Data

### Processing genomic data of SARS-CoV-2

In Dataset 1, 6000 sequences sampled from 6 regions (Africa, Asia, North America, Oceania, South America) in the period from 09/24/2020 to 09/25/2021 were downloaded from database NCBI Virus. In Dataset 2, we used 20,000 random sequences from the NCBI Virus database for the same period of time as in Dataset 1.

The sequences were not stratified by sampling date for two reasons, as follows. Firstly, the necessary size of statistics would be missing. Also, previous experience with hemagglutinin data [39] demonstrates that combining sequences from different dates is an advantage, because one compares earlier and later strains and thus finds a fitness valley connecting them. In contrast, estimation of selection coefficients, indeed, requires classification by time [70] .

Downloaded data were processed by a custom-made sequence-processing program *ProcessGenomes.py*. Multiple alignment was performed in MAFT online, using the “Fast calculation of full-length MSA of closely related viral genomes” [71]. From the total number of downloaded and aligned sequences, sequences in which the percentage of symbols “–“ (gap) or “N” (nonread) is greater than *Sequence threshold* are removed. From the remaining sequences, sites in which the percentage of gaps or letters N in across sequences is higher than *Gap threshold* are removed. Then, sites with a polymorphism degree (Hamming distance) lower than *Monomorphism threshold* are removed. We put the monomorphism threshold at two values (Table 1) and make sure that the choice does not affect final results, i.e., results are in a plateau in this parameter. The sequence data are binarized: the variant similar to consensus defined as Wuhan variant is declared 0, the others are 1, with the resulting population matrix *K_ni_*.

After binarization, the sequence data of virus undergo an additional processing stage, as follows. Based on the number of differences from sample consensus, a mutational spectrum is constructed using a custom-made sequence processing program *ProcessGenomes.py* (Figs. 3 and 5). Several groups of sequences are identified from this diagram. The new matrix *K_ni_* is formed by random sampling from two chosen groups, with the possibility of repetition.. The dependence of the predicted number of epistatic links on the sample size for simulated data is shown in Figs. 1 and 4.

### Epistasis detection and measurement

Based on matrix *K_ni_* obtained as described above, the universal footprint of epistasis [36] denoted *UFE_ij_* is calculated for each possible pair of sites *i*, *j* as 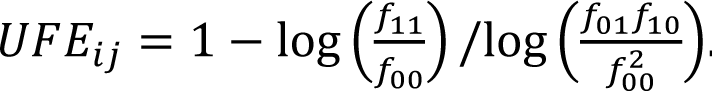 Then, the conditional coefficient *UFE_ij_*_0_ is re-calculated for each locus pair [39], as follows. Only sequences where at least one of the neighboring loci of the pair has allele 0 are used. The minimum of all UFE over such neighbors is denoted *UFE_ij_*_0_. Because the array of data used is large, these coefficients are calculated using parallel computing on the video card with more than 1000 parallel processors with the help of CUDA language. Pairs of loci, where the ratio of *UFE_ij_* to *UFE_ij_*_0_ is equal to or greater than 0.5, are considered “true pairs”, while the remaining pairs are excluded. Results are displayed in the form of circular diagrams (Figs. 6 and S5).

### Error calculation

The estimated epistatic pairs are compared with the matrix of true pairs *T_ij_* set in the Monte Carlo simulation code that generates genomic sequences for testing the accuracy. The relative error of the method is calculated as the sum of the number of false positive and false negative pairs for an initially–uniform population (Fig. 4). For the case of standing variation, the sum is divided by the total number of true pairs (Fig. 1).

### Assumptions and approximations

The method implies the weak selection limit, defined as max(1/*N*, μ) ≪ *s* ≪ 1, and is appropriate in the travelling wave regime, when many mutations at different loci are being fixed gradually and compete with each other for the space in population (clonal interference). The traveling wave regime takes place at *NμL* ≫ 1 and on time scales longer than 1/*s*. Next, the fluctuations of mutation rate along the genome, which are typically strong for RNA viruses, do not matter much in the multi-site evolution regime, where the mutation rate enters all the observables in the argument of a large logarithm [53].

Recombination, although observed for example for Omicron variants, is relatively rare in SARS-CoV-2, because it requires coinfection of an individual with two genetically remote variants, as estimated before for another virus [69, 72]. Multiplicity of infection is incorporated into an effective recombination rate measurable directly from genomic data [69, 73]. The effect of recombination on detection accuracy was tested an initially-diverse population and found to be modest (Fig. 1).

Also, here we consider the population level evolution only, because SARS–CoV-2 is acute and represented by a single variant in 99,5% patients, similar to influenza. In principle, the 0.5% chronic patients could seed new variants, in which case inter-scale evolutionary conflicts are possible [74].

## Supporting information

Reply to Reviewers

## Acknowledgements

We express our gratitude to Richard Neher for stimulating comments. We thank A.A. Makashov and S.V. Lagunov for technical help and discussion. We are grateful to D. A. Gorshkova, A. I. Pikhulya, E. I. Stroganova, and E. A. Vigovskaya for technical help.

## Ethical statement

No ethical statement is required.

## Availability of data and software

Codes are available at https://github.com/FireFrog17/Fundamental-restriction-on-epistasis-detection-and-fitness-valleys-in-virus-evolution-Data

## Supplementary material

**Figure S1.**
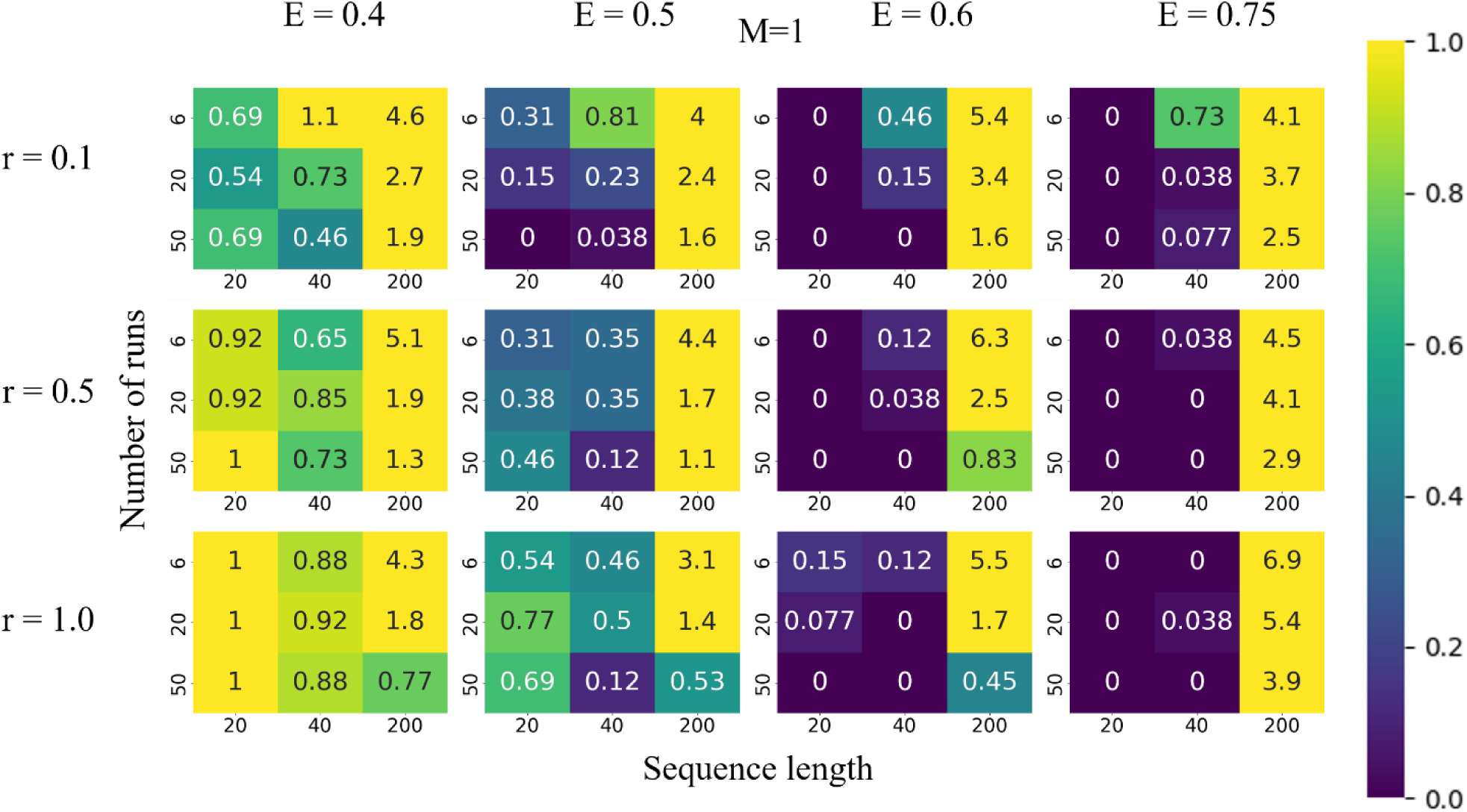
Accuracy of detection of epistatic pairs from UFE correlator for *M* = 1. Simulation parameters and notation are as in Figure 1.

**Figure S2.**
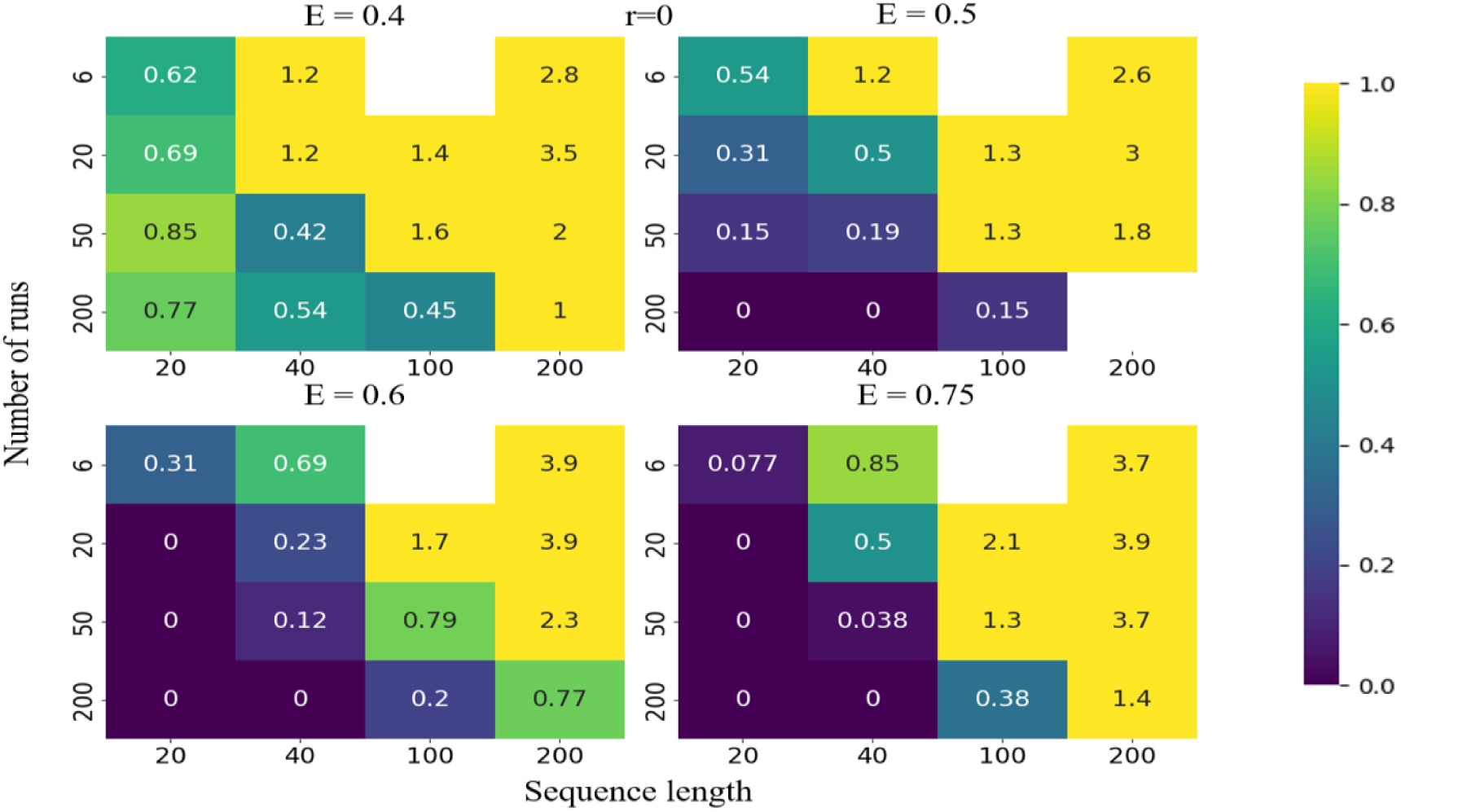
The error of detection of epistatic interactions from UFE correlator in the absence of recombination. Simulation parameters and notation as in Figure 1.

**Figure S3.**
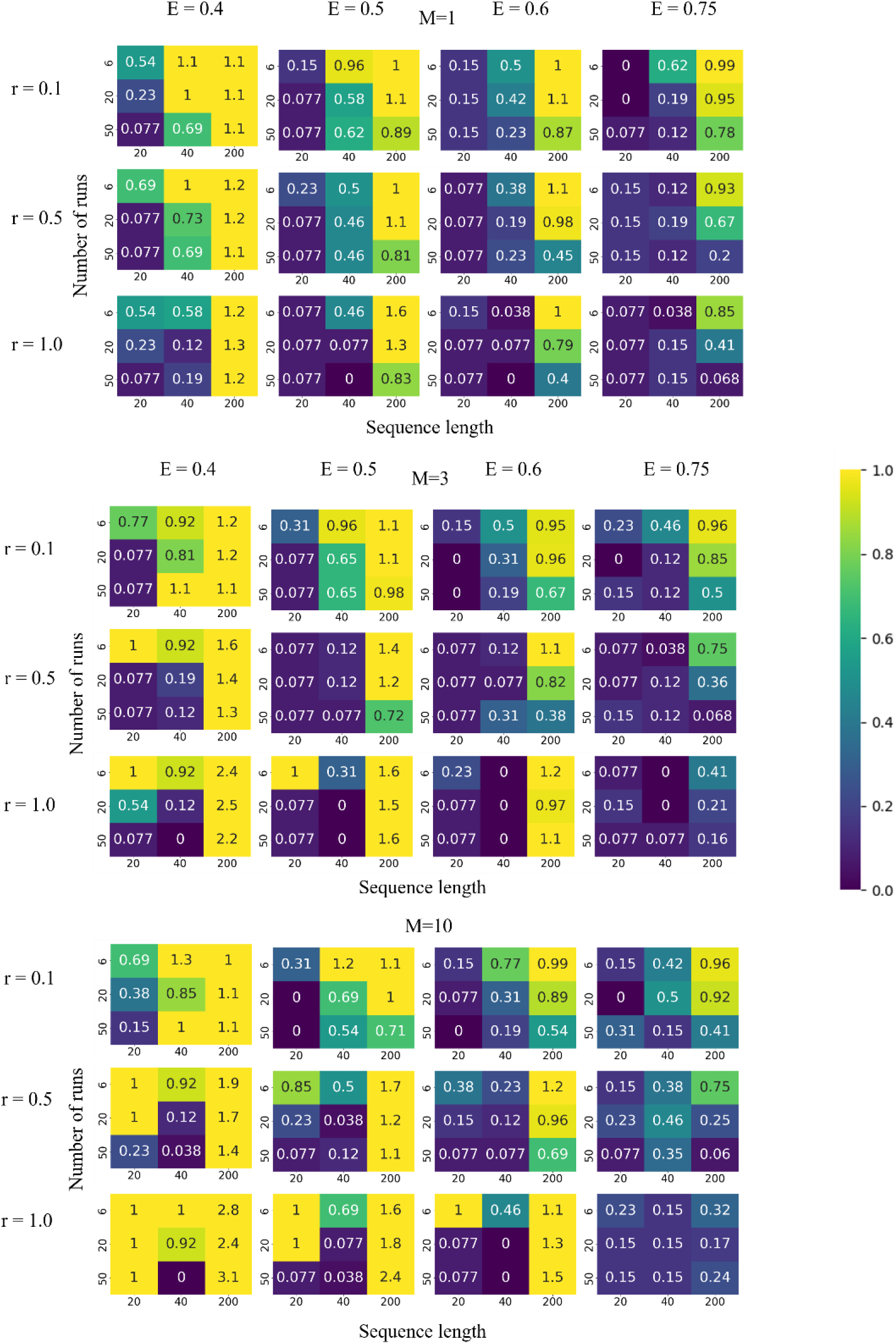
The error of detection of epistatic interactions from Pearson coefficient. Simulation parameters and notation as in Figure 1.

**Figure S4.**
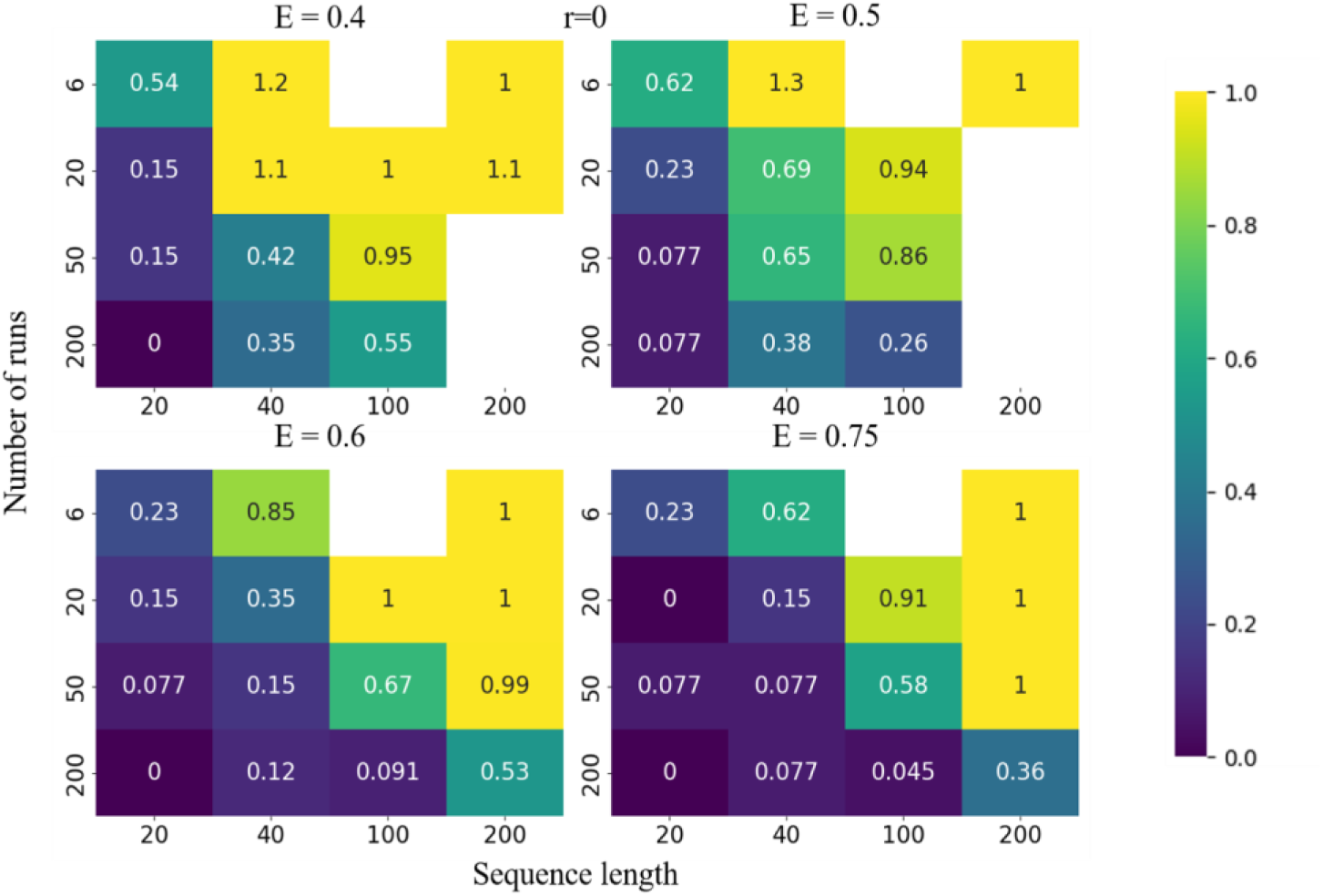
The error of detection of epistatic interactions in the absence of recombination from Pearson coefficient. Simulation parameters and notation are as in Figure 1.

**Figure S5.**
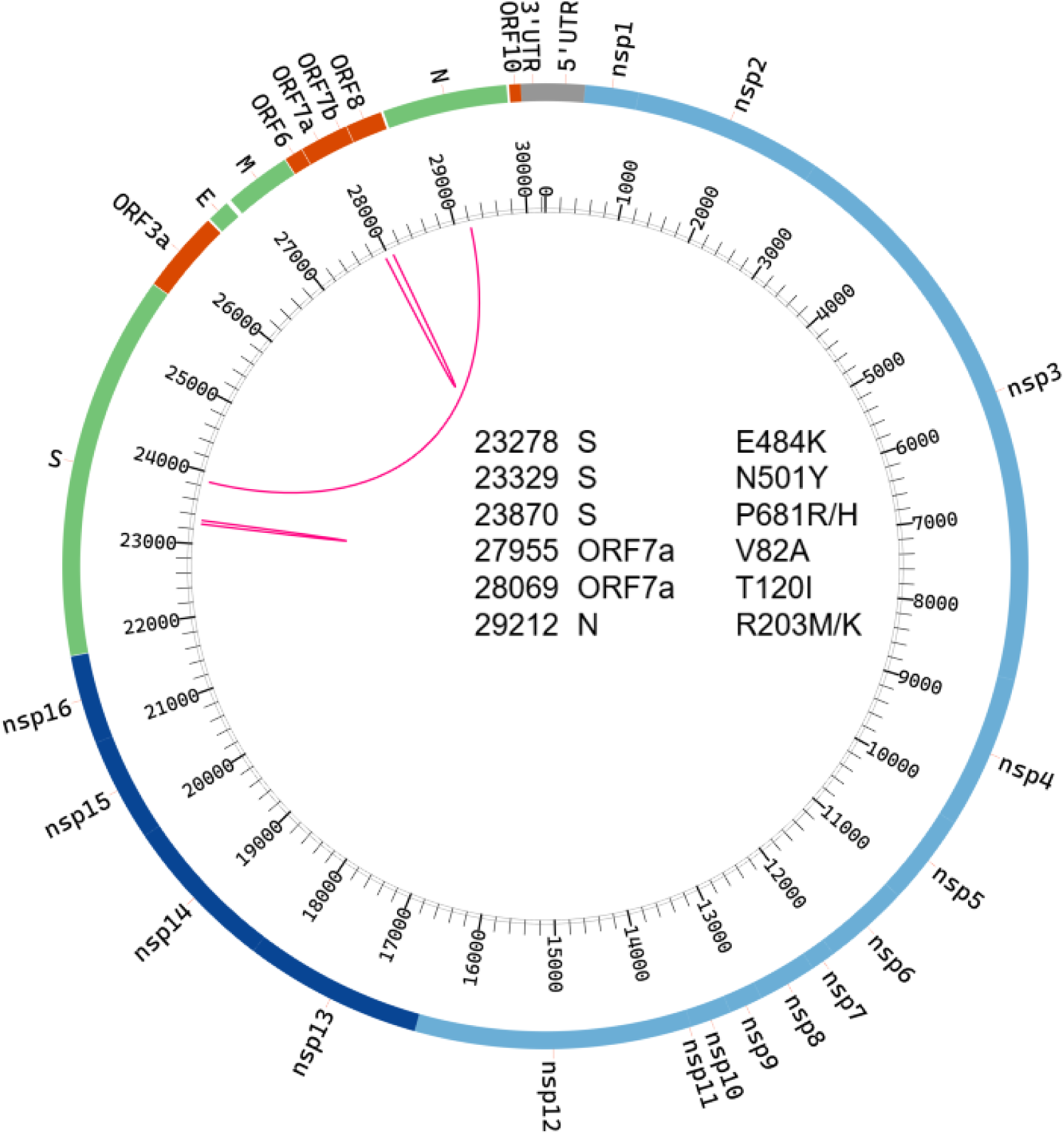
Epistatic interactions for SARS-CoV-2 genome predicted for Dataset 1 between groups I and III in Fig. 5. Notation as in Fig. 6. To obtain the circular diagram using software Circos [40], group I was oversampled 10 times and group III was oversampled 20 times, both with replacement.

**Figure S6.**
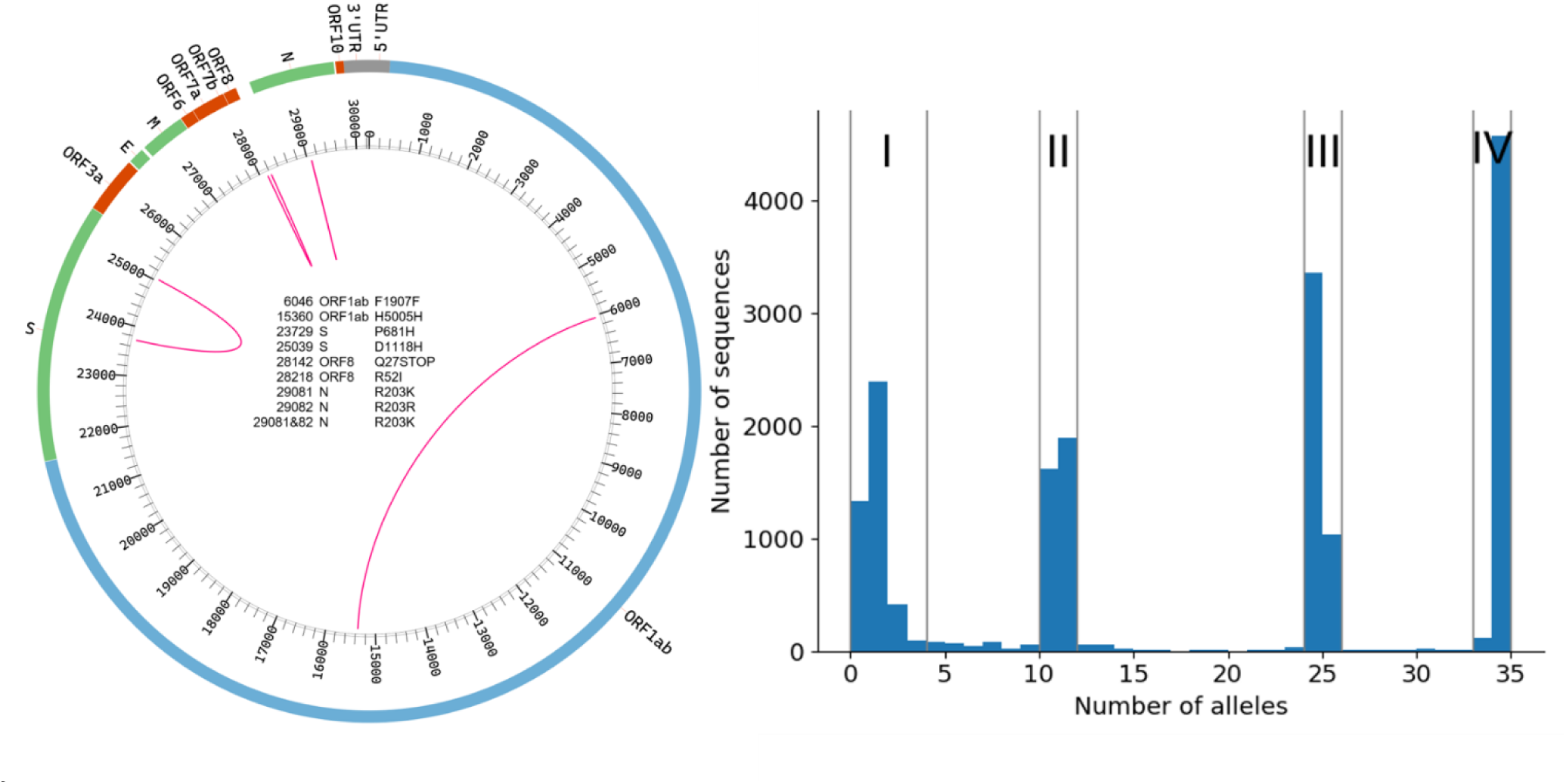
Epistatic interactions for SARS-CoV-2 genome predicted for Dataset 2 from UFE correlator. Dataset 2 was constructed using 20,000 random sequences from the NCBI without geographic classification. Virus database for the period from 09/24/2020 to 09/25/2021. Calculations were based on equal mixture of sequences from peak I and II.

