## Supplementary material for "Epistasis detection and fitness valleys in viral evolution": Reply to Reviewers

Line numbers given below correspond to the revised manuscript version with highlighted changes.

**Reviewer #1 :**

**Likhachev and Rouzine explore how accurately epistatic interaction can be measured from genetic diversity data using a method they previously developed. They first generate synthetic data and investigate how well the method discovers the ground truth and then apply the method to SARS-CoV-2 data.**

Thank you for describing the structure of our work correctly.

**If I understand correctly, what the authors refer to as mutation spectrum is essentially the distribution of Hamming distances from a consensus sequence.**

Yes.

**I have two points here: (i) why not compare to a reference sequence?**

In the original manuscript, we used the consensus of our Dataset 1 as the reference sequence. In the revised version, we use Wuhan variant as the reference sequence. We tested by simulation that taking the initial genomic sequence as the reference gives a higher detection accuracy.

**and (ii) there are many sequences that seem far too diverged for the indicated period ending in September 2021. The largest distances from the reference (or a consensus) in Sept 2021 should have been 50, maybe 60 mutations. It seems to me that the alignment is not very clean or that problematic sequences were included in the analysis.**

The focus of the manuscript is on mixing sequence groups I and II (Figs. 5 and 6), where the distance from Wuhan variant does not exceed 40 alleles. It is possible that sequences with large deletions or insertions are present in group III, where the distance is larger. When we mix I and III, we obtain a subset of links detected from groups I and II (Fig S5).

**The simulations seem rather unrealistic for the SARS-CoV-2 application. they are initialized as diverse population where every deleterious allele is at frequency 0.45. SARS-CoV-2 started as a monomorphic outbreak and most sites have very low allele frequencies. Furthermore, results in Fig 1 explore recombination rates of 0.1, 0.5, and 1.0. SARS-CoV-2 recombination rate depend on co-infection and are likely much lower. Linkage is thus a much more serious problem...**

To address this important concern, we added a new section (Figures 2-4), where we test the detection accuracy of the method for the initially–monomorphic population with no recombination. To explain the emergence of VOC, we also assume positive epistasis in the form of a fitness valley (the primary mutation and 10 compensatory mutations, Fig. 2 insert). As expected, the detection error is much higher for an initially monomorphic population than for an initially polymorphic population (cf. Fig 4 and Fig 1), but it does not cancel the utility of the method. The technique misses a half of compensatory sites, the weaker half, but on the bright side it does not generate any false positives (Fig. 4 and the legend).

**... and most previous approaches have used independent mutation events for fitness estimation.**

We do not address the fitness effects of independent mutations here, because it requires a separate method. Such estimation for E Coli is addressed here:
https://www.biorxiv.org/content/10.1101/2025.11.26.690712v1

**Is there any biological context or rationalization for the interacting pairs flagged by the method?**

Yes, there is. We added a paragraph in Results (Lines 300-320):

Mutations E484K and N501Y are well-known mutations of the receptor-binding domain of the coronavirus in Spike. The first is responsible for the viral escape from the immune response and is capable of reducing the affinity of antibodies produced to earlier strains by 1-2 orders of magnitude, while the second improves the binding of coronavirus to the human ACE2 receptor on cell surface and increases viral replication in upper respiratory tract cells. [41]

Mutations P681H and P681R, most often found in Alpha (B.1.1.7) and Delta (B.1.617.2) variants of the virus, respectively, are located in the furin cleavage region of Spike. They both confer a change in the amino acids from nonpolar hydrophobic to positively charge. They affect tropism (repertoire of infectable cells) by increasing the cleavage of S1/S2 proteins in human airway epithelial cells [42].

Mutation D138Y in Spike is believed to affect the protein's secondary structure thus reducing antibody - binding efficiency [43]. The R203K mutation in the N protein was shown to increase transmission and virulence of SARS-COV-2 [44], while the R203M mutation, which creates a start codon for a new open reading frame, was reported to suppress interferon induction, thereby increasing viral infectivity [45].

V82A and T120I in ORF7a are most frequently found in variant Delta of the coronavirus [46]. They are located within the di-lysine motif of ORF7a, which is important for the localization of viral proteins in endoplasmic reticulum. Existing data on mutation Q7036H in protein ORF1ab and mutation V71I in ORF7a do not permit to determine their functions definitively; however, their location within viral non-structural proteins hint at their interaction with the immune system of the host.

**It would be much preferable to list these as amino acid substitutions rather than nucleotide positions.**

Done (Figs. 6 and S5).

**While I agree that the application of QLE based methods to SARS-CoV-2 is ill-advised and that the inherent assumptions are strongly violated…**

Thank you for agreeing with this important point.

**… I don't think the authors have convincingly shown that their approach does a lot better.**

We disagree. Our paper tests for the accuracy and does not neglect linkage, so we do a lot better. The advantages of our approach are now explained in Introduction, as follows (Line 111):

In the present work, we test the fidelity of this detection technique using Monte-Carlo simulation in a broad range of parameters. Then, we apply the method to real virus sequences from the adapting world population of SARS–CoV-2 sampled in two different ways to predict the strongest epistatic links. This approach differs from the alternative methods in the following respects. Firstly, it does not make unrealistic assumptions regarding the evolutionary factors at play. Secondly, its accuracy is validated in a broad range of model parameters on simulated sequences, using a generic model of population genetics of the Wright Fisher type. We demonstrate the accuracy on simulated sequences under conditions resembling the evolution of SARS CoV-2 including the emergence of variants of concern. Thirdly, the application to data is focused on well-balanced sampling between sequences related to major variants. The fourth feature is the use of the conditional correlator.

**In fact, tree based methods (like Neverov et al) take strong linkage effect into account explicitly.**

In fact, the use of a phylogenetic tree does not remove linkage effects in epistatic detection (Pedruzzi & Rouzine PLoS ONE 2019). We added to Discussion (Line 428):

Note that the use of tree-based methods does not automatically compensate for strong linkage effects. This is because, in the absence of sufficiently strong recombination, all sequences in one single population are very similar to their MRCA, and MRCAs for different independent populations are diverging from each other in genetic space. This process was demonstrated to be responsible for strong linkage effects [34]. To decrease the effects of linkage, one group used synonymous sites as markers of linkage [37, 38].

**These can be applied to very large datasets and should extract all available information from the data.**

The fact that the same work can be done by another method is hardly a valid critique. We agree that it would be interesting to compare the two approaches, which we plan to do in the future. Note that the potential future work proposed by Richard must use balanced averaging over data and testing by simulation, as we do in our work.

**Minor points:**

**- why not break down the sequence by variant or Pango lineage? The Hamming distance distributions should be readily interpretable in terms of pairwise comparison between variants.**

As stated in the title of the manuscript, we seek fitness valleys connecting major variants to explain the emergence of VOC. Hence, we use the balanced mix of variants isolated from different time points.

**- QLE does not ignore linkage effects. It assumes that recombination is rapid compared to allele frequency changes.**

In the end, it amounts to the same thing: linkage effects are negligible. We edited the sentence (line 394):

"quasi linkage equilibrium" approach [28, 30], which assumes frequent recombination compared to changes in allelic frequency and a large population size. In this limit, genetic linkage effects are negligible compared to epistasis and recombination.

**Signed: Richard Neher**

Thank you for very stimulating comments.

**Reviewer #2 :**

**This paper uses an epistasis inference method (developed by Pedruzzi & Rouzine 2021: [https://journals.plos.org/plospathogens/article?id=10.1371/journal.ppat.1009669](https://journals.plos.org/plospathogens/article?id=10.1371/journal.ppat.1009669)) to understand how the accuracy of detection (i.e., a normalized sum of false positives and negatives) depends on features such as sequence length. The method is suggested as an alternative to methods of epistasis inference based on patterns of covariance between mutations (i.e., based on linkage (dis)equilibrium; [https://www.pnas.org/doi/full/10.1073/pnas.2012331117](https://www.pnas.org/doi/full/10.1073/pnas.2012331117)) and phylogeny (i.e., the nature of substitutions occurring along the tree; [https://journals.plos.org/plosgenetics/article?id=10.1371/journal.pgen.1001301](https://journals.plos.org/plosgenetics/article?id=10.1371/journal.pgen.1001301)). The key steps are to calculate some measure of covariance or correlation (e.g., r2) for SNPs at two loci, test the significance of the association and retain SNP pairs that are sufficiently correlated, and calculate *conditional* correlation coefficients (given "a locus connected to the pair of interest carries the consensus allele...**

Thank you for the correct description of our procedure.

**which I take to mean "of the ancestral type").**

In the revised version, we use Wuhan variant as the consensus.

**The authors test this method using simulations to test the ability of their method to recover epistasis pairs and, applying their method to SARS-CoV-2 data, identify such interactions.**

**I think the core idea of the method is interesting…**

Thank you for the positive opinion about our core idea.

**… but the paper in its current form is difficult to read and the logic of the method, simulations, and inferences drawn from the SARS-CoV-2 data are underdeveloped. The paper would benefit by carefully explaining the approach…**
**… and ensuring that confounding in the empirical portion of the paper is controlled for in some capacity.**

In the revised version, we did our best to explain our approach based on Reviewers' comments.

**… appropriately benchmarking it to LD- and tree-based methods, putting the study firmly in the context of previous work on epistasis and SARS-CoV-2 evolution…**

In the revised Discussion, we discuss these methods in more detail. Our final results are compared to the results of two other methods in Fig. 7.

**… 5. In the current version of the paper, all comparisons with other methods of pairwise epistasis inference (eg, those that use substation histories on trees) are strictly verbal. In particular, there is no evidence based on the simulations that other methods to epistasis detection fare poorly in any part of parameter space (i.e., as a function of mutation rate, genome size, selection strength, etc.). I could not find comparisons in the papers on which this study builds, either.**

The problem is that their authors usually do not test their accuracy on simulated sequences that evolved under known conditions with preset epistatic parameters, which can be then compared to the inferred values. One contribution of the present work is to use such testing as a standard routine (Figs. 1 and 4).

We also compare the final results for SARS CoV 2 between three methods (Fig. 7).

**Such comparisons are necessary to conclude "fundamental limitations" on epistasis detection exist and to properly contextualize the method in terms of what has come before.**

Fundamental limitations are called "fundamental", because they do not depend on a detection method used, but are intrinsic to evolutionary dynamics. The existence of strong correlations caused by genetic linkage is known in population genetics for more than 100 years. To make it clear, we added a paragraph to Introduction (Line 84):

About a century ago, it was realized that the evolution of a population is strongly affected by the fact that the fates of alleles at different loci are linked unless separated by recombination [33]. Empirically, the effect of strong linkage between genomic sites has been discovered by Thomas Hunt Morgan who used the inter-dependent segregation of alleles in fruit flies to understand the role of chromosomes and sex in inheritance. Since then, a large body of experiment and modeling demonstrated that linkage changes evolutionary properties drastically. In the absence of recombination, common phylogeny makes all sequences very similar to their

MRCA [34]. Moreover, even very frequent recombination does not break up linkage effects in a sufficiently long genome [35].

and below (Line 111):

In the present work, we test the fidelity of this detection technique using Monte-Carlo simulation in a broad range of parameters. Then, we apply the method to real virus sequences from the adapting world population of SARS–CoV-2 sampled in two different ways to predict the strongest epistatic links. This approach differs from the alternative methods in the following respects. Firstly, it does not make unrealistic assumptions regarding the evolutionary factors at play. Secondly, its accuracy is validated in a broad range of model parameters on simulated sequences, using a generic model of population genetics of the Wright Fisher type. Thirdly, the application to data is focused on well-balanced sampling between sequences related to major variants. The fourth feature is the use of the conditional correlator.

**Major comments:**

**1. The description of the method ("The method of detection of epistatic pairs upgraded from (39) consists of three stages, as follows. (i) Based on the binarized sequence data, a paired correlator, such as UFE (35) and Pearson coefficient, is calculated for each pair of genetically-diverse genomic sites (loci). ... Based on the comparison between these correlators, one determines whether the candidate pair is epistatic or not, as described in Methods.") is insufficient and understanding it, even in broad strokes, requires diving into previous papers of the authors.**

In the revised version, we made a better description of the method in Methods (Line 505):

**Epistasis detection and measurement.** Based on matrix $K_{ni}$ obtained as described above, the universal footprint of epistasis [36] denoted $UFE_{ij}$ is calculated for each possible pair of sites $i, j$ as $UFE_{ij} = 1 - \log\left(\frac{f_{11}}{f_{00}}\right) / \log\left(\frac{f_{01}f_{10}}{f_{00}^2}\right)$. Then, the conditional coefficient $UFE_{ij0}$ is re-calculated for each locus pair [39], as follows. Only sequences where at least one of the neighboring loci of the pair has allele 0 are used. The minimum of all UFE over such neighbors is denoted $UFE_{ij0}$. Because the array of data used is large, these coefficients are calculated using parallel computing on the video card with more than 1000 parallel processors with the help of CUDA language. Pairs of loci, where the ratio of $UFE_{ij}$ to $UFE_{ij0}$ is equal to or greater than 0.5, are considered "true pairs", while the remaining pairs are excluded. Results are displayed in the form of circular diagrams (Figs 6 and 7).

**This is a problem because there are important limitations to the method. Key among the limitations of the method is that the three sites in the conditional correlation calculation must be polymorphic. This places strong restrictions on the precise contexts in which the statistics can be meaningfully calculated and interpreted.**

All methods of epistasis detection consider only polymorphic sites. Well-conserved sites carry no information in this regard.

**It also makes tuning hyper-parameters (e.g., which sequences to include in the analysis) very challenging.**

This subtle point is explained in the revised version (Line 493):

We put the monomorphism threshold at two values (Table 1) and made sure that the choice does not affect final results, i.e., results are in a plateau in this parameter.

**If mutations are instantaneously fixed (i.e., selection is much stronger than mutation) it would be inappropriate to calculate the conditional correlation coefficient.**

This scenario is not very realistic for an RNA virus. Mutations with a large selection coefficient s ~ 1 that are fixed instantaneously (within 1-2 generations) are extremely rare in any real system including RNA viruses. Even if selection coefficient is much stronger than mutation rate, its value is usually much smaller than unit. All such mutations are fixed very gradually, with the half-time of fixation inversely proportional to the selection coefficient and logarithm of the ratio of selection coefficient to the mutation rate (Haldane, 1927). Hence, the fixation times are large. Linkage effects, which are always strong for RNA viruses including those with recombination, make the fixation process even slower. For example, in a drug-treated HIV patient, it takes 4 weeks to half-fix a primary ritonavir resistance mutation. In the case of SARS-CoV-2 at the population level considered here, the time scales are months.

**I can imagine that, in the weak selection limit, the method is appropriate.**

The method works if selection coefficient is much stronger than mutation rate, but both are very small compared to unit, This situation is typical for rapidly–evolving RNA viruses. We added this condition to Lines 520-521:

The method implies the weak selection limit, defined as $\max(1/N,\ \mu) \ll s \ll 1$….

**But it is unclear to me if this situation is similar to the quasi-linkage equilibrium regime, in that LD will have broken down via recurrent mutation (given enough time).**

The real situation with SARS –CoV-2 and our simulation model are very different from QLE limit, which neglects linkage effects between numerous sites, assuming that recombination is very strong and the population size is enormous. QLE method is discussed in Discussion (Lines 394-403). We added in the revised version (Lines 425-227):

QLE does not work in SARS CoV-2 evolution, because linkage effects, which are negligible in QLE limit, are strong. Lastly, a population of infected individuals is not as large as required for QLE to work.

**What are the relevant timescales and regime boundaries?**

Added in Methods/Assumptions and approximations:

The method implies the weak selection limit, defined as $\max(1/N,\ \mu) \ll s \ll 1$, and is appropriate in the travelling wave regime, when many mutations at different loci are being fixed gradually and compete with each other for the space in population (clonal interference). The traveling wave regime takes place at $N\mu L \gg 1$ and on time scales longer than $1/s$.

**2. Does the analysis of the SARS-CoV-2 control (in any way) for violations of the method's assumptions (e.g., hyper-mutable or recombinagenic parts of the genome)? It appears not… …Moreover, there is now ample evidence that particular regions of the genome are more likely to exhibit polymorphism than others due to intrinsic and extrinsic differences in mutation and recombination.**

Explained in Methods/Assumptions (Line 524):

Next, the fluctuations of mutation rate along the genome, which are typically strong for RNA viruses, do not matter much in the multi-site evolution regime, where the mutation rate enters all the observables in the argument of a large logarithm [53].

Recombination is relatively rare in SARS-CoV-2, because it requires coinfection of an individual with two genetically remote variants, as estimated before for another virus [69, 72]. Multiplicity of infection is incorporated into an effective recombination rate measurable directly from genomic data [69, 73].

**I could not find evidence that sequences are stratified by sampling date even though this will determine the amount of polymorphism observed . and lead to problems in the calculation of**

**the unconditional and conditional correlation coefficients. Controlling for the issues of timescale and evolutionary rate heterogeneity are essential in epistasis inference.**

It depends on the aim of a study. The aim of the present study is to detect the fitness valley related to VOC emergence. That task requires mixing sequences from different time points, before and after VOC emergence. Explained in Methods (Line 481):

The sequences were not stratified by sampling date for two reasons, as follows. Firstly, the necessary size of statistics would be missing. Also, previous experience with hemagglutinin data [39] demonstrates that combining sequences from different dates is an advantage, because one compares earlier and later strains and thus finds a fitness valley connecting them. In contrast, estimation of selection coefficients, indeed, requires classification by time [70] .

3. The authors aim to identify the *signatures* of epistasis, but epistasis is always defined with respect to a phenotype (e.g., fitness, the binding affinity to sialic acid receptors in a particular context). It is not clear, in the data analysis, what kind of epistatic interactions are being detected. To the best of my knowledge, it is a kind of time-averaged epistasis in within- and between-host fitness.

Added in Intro (Lines 76-77):

…where epistasis is defined with respect to fitness, i.e. the reproduction number at the within-host or between-host levels .

**There may also be some sex-, host-, and space-averaging going on in the background, too, depending on the exact application and the tuning of hyperparameters. It is difficult to say, which in turn makes interpreting the data and making comparisons challenging.**

Added in Methods/Assumptions (Line 533):

Also, here we consider the population level evolution only, because SARS–CoV-2 is acute and represented by a single variant in 99,5% patients, similar to influenza. In principle, the 0.5% chronic patients could seed new variants, in which case inter-scale evolutionary conflicts are possible [74].

**4. I am not convinced that the simulations are appropriate.**
**4a- The Wright-Fisher model with recombination is insufficient to describe the evolutionary dynamics of systems such as SARS-CoV-2 and influenza, which the authors are interested in describing. A multi-locus epidemiological model is more appropriate, and formatting such a model requires specifying how mutations affect various, correlated components of fitness (replication, transmission, virulence). For a number of phenotypes, the conditional correlation coefficient which is used to infer epistasis would likely capture the effective epistasis in total fitness over a period of time. The WF model is almost certainly misspecified, and unfortunately it is difficult to determine *how* the misspecification affects the conclusions of the simulation study portion of the paper. I recognize it is impractical to adjust the paper to reflect issues in this respect, but justification should be given for in what ways the WF model is or is not appropriate.**

The multi-locus epidemiological model has been shown to map onto a WF model with an effective selection coefficient distribution by three modeling teams. This point is explained now in Discussion (Line 380):

For testing the accuracy, we used a model with directional selection. In fact, SARS-CoV–2 evolves in antibody–binding regions due to collective immunity of population, which is a case of the selection for change (Red Queen effect). Nevertheless, as it has been shown by several groups [56-58], the multi-locus epidemiological model of respiratory viruses is effectively reduced to the Wright-Fisher model with constant directional selection.

The selection coefficient is expressed in terms of basic reproduction ratio in a naïve population and the breadth of antibody neutralization, calculated in these works using different approximations.

**4b- Viruses range from clonal to highly recombinagenic. Recombination also scales with the density of infections in a non-linear way. This is not reflected in the treatment of recombination in the simulations and I would not be surprised if the "crossover" frequency parameter is such that the systems simulated are closer to the quasi-linkage equilibrium attractor over a substantial portion of the parameter space (for values of E not close to 1).**

We now explained in Methods/Assumptions why the role of recombination for detection of epistasis is small (Line 527):

> Recombination, although observed for example for Omicron variants, is relatively rare in SARS-CoV-2, because it requires coinfection of an individual with two genetically remote variants, as estimated before for another virus [69, 72]. Multiplicity of infection is incorporated into an effective recombination rate measurable directly from genomic data [69, 73]. The effect of recombination on detection accuracy was tested an initially-diverse population and found to be modest (Fig. 1).

**I also have more minor quibbles with the figures, e.g., the suppression of values of the "error detection" statistic. Why not make figures showing how Type I and II error scale with sequence length and replicate population sizes?**

In the legend of Fig. 1, the error of detection is defined as the sum of the false positive and false negative interactions divided by the number of true interactions averaged over populations whose number is shown. This is the relevant measure of the accuracy of detection.

**Moreover, it is not clear to me if the values used for sequence length are appropriate. Does the method not scale well when the genome length is >200 (presumably, in units of base pairs)?**

Unfortunately, we cannot run simulations at lengths much larger than 200, as they take too much time.

**Overall, I think this method of epistasis inference is useful in certain regimes if those regimes can be mapped out, limitations explored and discussed, and the links to previous work made explicit.**

We hope that our revisions helped to clarify these issues.

**I think clarity about what information is being utilized and in what manner (in the method and data analysis) is essential, and recommend that the authors consider the meaning of the epistatic pairs that are being inferred as well as the kinds of model structures (WF or otherwise) to which the method is being applied in silico.**

Thank you for the comments.

**Reviewer #3:**

**This is an additional contribution in a series of papers on the Universal Footprint of Epistasis framework, a method that attempts to detect epistasis between sites using segregating variation. In general, random linkage effects result in correlations between sites that are not due to epistasis, and the Universal Footprint of Epistasis attempts to control for this effect. The current manuscript first applies this framework in the context of a simulation study to determine conditions when epistasis can be reliably detected using this method. It then applies the Universal Footprint of Epistasis framework to genome-wide SARS-CoV2 sequences, detecting a**

**small set of interacting mutations.**

**The title of the piece is "Fundamental restriction on epistasis detection and fitness valleys in virus evolution", however I do not think that the readers of GENETICS will feel that there is a fundamental result presented.**

In the revised version, we changed the title to:

"Epistasis detection and fitness valleys in viral evolution"

**The paper assumes a great deal of familiarity from the reader about the Universal Footprint of Epistasis framework and does not present the main test being used, and so the paper is inaccessible to a general readership.**

We revised the work to make it more accessible.

**At the same time, the simulation study, which presumably is the evidence for the fundamental restriction....**

Fundamental restriction was shown previously, starting from Morgan (Line 77):

About a century ago, it was realized that the evolution of a population is strongly affected by the fact that the fates of alleles at different loci are linked unless separated by recombination [33]. Empirically, the effect of strong linkage between genomic sites has been discovered by Thomas Hunt Morgan who used the inter-dependent segregation of alleles in fruit flies to understand the role of chromosomes and sex in inheritance. Since then, a large body of experiment and modeling demonstrated that linkage changes evolutionary properties drastically. In the absence of recombination, common phylogeny makes all sequences very similar to their MRCA [34]. Moreover, even very frequent recombination does not break up linkage effects in a sufficiently long genome [35].

The increasingly dominant effect of linkage over epistasis in allelic associations originates from the genetic divergence of independent populations in time. All sequences in a population are similar to their most recent common ancestor, which moves away from the origin along a stochastic trajectory [34]. As a result, any measure of co-variance produces strong noise of random sign. An attempt at detecting epistasis, whether by using covariance measures, such as Pearson coefficient, mutual entropy, universal footprint [36], or by tree-based methods [37, 38] faces the same problem, the overwhelming genetic linkage effects. The only way to resolve this issue is to average the haplotype frequencies over many independent populations evolving under similar conditions. To decrease the required number of populations, it was proposed to use only sequences with a majority allele at a neighbor site of the measured site pair to interrupt a path along sites that creates a false-positive interaction [39].

The aim of the present paper is to test a working method of detection, which that takes this well–known fact into account, on simulated sequences, in a broad range of parameters (Line 88):

In the present work, we test the fidelity of this detection technique using Monte-Carlo simulation in a broad range of parameters. Then, we apply the method to real virus sequences from the adapting world population of SARS–CoV-2 sampled in two different ways to predict the strongest epistatic links.

**...., looks at a rather specific set of scenarios and thus is lacking generality to support the title claim. .... I also did not find it to add very much beyond the previous documentation in this series of papers on the difficulty in detecting epistatic interactions from patterns of segregating variation in a manner that is grounded in population genetic theory.**

PREVIOUSLY my team investigated a single parameter set and a single evolutionary scenario with a uniform periodic epistatic network. The NOVEL contribution of Igor Likhachev is to investigate the accuracy of the method **in a broad range of parameters.** Also, we now added a new evolutionary scenario more representative of SARS CoV-2, as proposed by Richard Neher (Reviewer 1).

PREVIOUSLY we applied it to influenza sequences. NOVEL contribution of Igor Likhachev is to apply it to SARS CoV 2 sequences.

Novelty is stated in Introduction, as follows:

In the present work, we test the fidelity of this detection technique using Monte-Carlo simulation in a broad range of parameters. Then, we apply the method to real virus sequences from the adapting world population of SARS–CoV-2 sampled in two different ways to predict the strongest epistatic links. This approach differs from the alternative methods in the following respects. Firstly, it does not make unrealistic assumptions regarding the evolutionary factors at play. Secondly, its accuracy is validated in a broad range of model parameters on simulated sequences, using a generic model of population genetics of the Wright Fisher type for two scenarios of evolution. We demonstrate the accuracy on simulated sequences under conditions resembling the evolution of SARS CoV-2 including the emergence of variants of concern. Thirdly, the application to data is focused on well-balanced sampling between sequences related to major variants. The fourth feature is the use of the conditional correlator.

**In terms of the empirical analysis, it is difficult to judge the results. The pattern of inferred interactions seems to depend in a sensitive manner on how the dataset to be analyzed and is composed...**

It does, and this is not a problem with the method, but rather a general projection. The dependence on the sample composition with respect to geography structure (balanced), time, and reference sequence is another major contribution of the present work. We show that e well-balanced sampling between strains gives very stable results (Fig. 4)

**...and the results are also quite different from two different analyses of epistasis in SARS-CoV2 that used direct coupling analysis (DCA). These prior studies are described in this manuscript as being based on quasi-linkage equilibrium (QLE), but I am not sure the original authors would agree with this. Although both models express genotype frequencies as a quadratic function of indicator variables on the presence of allelic variants, QLE is based on an evolutionary model...**

I read the description of DCA in Rodriguez-Rivas et al and this is a version of QLE (see Neher and Shraiman review). We added in Discussion (Line 425):

…QLE does not work in SARS CoV-2 evolution, because linkage effects, which are negligible in QLE limit, are strong. Lastly, a population of infected individuals is not as large as required for QLE to work.

**whereas DCA is a phenomenological modeling approach. This difference leads to differences in practice, for example the Rodriguez-Rivas et al. PNAS paper down-weights sequences with over 80% identity, which is standard in the DCA literature (as a crude attempt to deal with the similar effects of random linkage as Universal Footprint of Epistasis) but would not make sense from a QLE standpoint.**
**… I agree with the authors that the practical success of DCA is perplexing from a population genetic perspective.**

We added a paragraph about DCA to Discussion (Lines 404-427):

To address the problem of false-negative bonds, another team [67] compared the sequences of SARS CoV-2 not between themselves, but with other coronaviruses. They used "direct coupling analysis", which postulates a statistical distribution of sequences (usually, of a Gaussian form) including individual effects of alleles and their pairwise interaction. A physicist would recognize here the Gibbs distribution for the Onsager model of ferromagnetism. In population genetics, this Gaussian form has a justification in the limit of QLE [28, 30]. Parameters of the Gaussian are fit to sequence data using various approximations [67]. DCA also includes down-weighing sequences with over 80% identity, which is a phenomenological trick to deal with the effects of random linkage. As result, the number of false-positive epistatic bonds due to genetic linkage is sharply decreased (Fig. 8C). The problem here is that the nature of such correction is not derived from a model of population genetics. QLE does not work in SARS CoV-2 evolution, because linkage effects, negligible in QLE limit, are quite strong.

**In any case, it is hard to evaluate the evidence for the claimed interactions because the manuscript does not contain a biological/functional analysis of the meaning and reasonableness of the inferred interactions...**

Thank you, this is very important. In the revised version, we added a paragraph devoted to the biological significance of the results (Lines 300-320):

Mutations E484K and N501Y are well-known mutations of the receptor-binding domain of the coronavirus in Spike. The first is responsible for the viral escape from the immune response and is capable of reducing the affinity of antibodies produced to earlier strains by 1-2 orders of magnitude , while the second improves the binding of coronavirus to the human ACE2 receptor on cell surface and increases viral replication in upper respiratory tract cells. [41]

Mutations P681H and P681R, most often found in Alpha (B.1.1.7) and Delta (B.1.617.2) variants of the virus, respectively, are located in the furin cleavage region of Spike. They both confer a change in the amino acids from nonpolar hydrophobic to positively charge. They affect tropism (repertoire of infectable cells) by increasing the cleavage of S1/S2 proteins in human airway epithelial cells [42].

Mutation D138Y in Spike is believed to affect the protein's secondary structure thus reducing antibody - binding efficiency [43]. The R203K mutation in the N protein was shown to increase transmission and virulence of SARS-COV-2 [44], while the R203M mutation, which creates a start codon for a new open reading frame, was reported to suppress interferon induction, thereby increasing viral infectivity [45].

V82A and T120I in ORF7a are most frequently found in variant Delta of the coronavirus [46]. They are located within the di-lysine motif of ORF7a, which is important for the localization of viral proteins in endoplasmic reticulum. Existing data on mutation Q7036H in protein ORF1ab and mutation V71I in ORF7a do not permit to determine their functions definitively; however, their location within viral non-structural proteins hint at their interaction with the immune system of the host.

**...or the nature of the disagreements with prior studies.**

We hope that, with all these new clarifications and additions, the nature of disagreement has been explained.

**Minor comments:**

**- Figure 5D should probably be redrawn since the graphic is from a previous study, and it is important that the authors have determined the interactions inferred from that study for comparison with their own.**

Done (now Fig. 8).

**- There is language used describing "bonds" in fitness valleys that I did not understand.**

Clarified: "(interactions)".

**- UFE should certainly be described at first use in the results.**

In the revised version, we refer the reader to Methods (Line 505):

**Epistasis detection and measurement.** Based on matrix $K_{ni}$ obtained as described above, the universal footprint of epistasis [36] denoted $UFE_{ij}$ is calculated for each possible pair of sites $i,j$ as $UFE_{ij} = 1 - \log\left(\frac{f_{11}}{f_{00}}\right) / \log\left(\frac{f_{01}f_{10}}{f_{00}^2}\right)$. Then, the conditional coefficient $UFE_{ij0}$ is re-calculated for each locus pair [39], as follows. Only sequences where at least one of the neighboring loci of the pair has allele 0 are used. The minimum of all UFE over such neighbors is denoted $UFE_{ij0}$. Because the array of data used is large, these coefficients are calculated using parallel computing on the video card with more than 1000 parallel processors with the help of CUDA language. Pairs of loci, where the ratio of $UFE_{ij}$ to $UFE_{ij0}$ is equal to or greater than 0.5, are considered "true pairs", while the remaining pairs are excluded. Results are displayed in the form of circular diagrams (Figs 6 and 7).

Thank you for the insightful comments.
